# Suppression of bacterial cell death underlies the antagonistic interaction between ciprofloxacin and tetracycline in *Escherichia coli*

**DOI:** 10.1101/2024.04.18.590101

**Authors:** James Broughton, Achille Fraisse, Meriem El Karoui

## Abstract

Antibiotic combinations are an attractive strategy to maximise the efficiency of drug treatment and minimise resistance evolution, but we still lack a full understanding of their effect on bacterial cells. The interaction between DNA-targeting antibiotics, such as ciprofloxacin, and translation inhibitors, such as tetracycline, is antagonistic, resulting in a weaker effect on bacterial growth than expected from the effect of each single drug. This antagonism has been analysed in detail at the population level, but we lack a single-cell understanding of its effect and how it depends on nutrient availability. Here, we used a microfluidic device to quantify the antagonism between ciprofloxacin and tetracycline at the single-cell level in three nutrient conditions. We showed that improved growth is due to increased survival of cells under the drug combination compared to ciprofloxacin alone. This effect is growth-dependent, with better suppression in rich nutrient conditions. Quantification of the DNA damage response (SOS response) revealed two sub-populations among the cells that die upon ciprofloxacin treatment, with some cells reaching a very high level of SOS while others had a lower level of SOS, similar to surviving cells. The low-SOS cells were more frequent in fast growth conditions and showed increased survival under the drug combination but the high-SOS cells were hardly rescued by the drug combination. This result explains the stronger antagonistic effect of tetracycline on ciprofloxacin in fast growth compared to slow growth. Our results underscore the importance of single-cell quantification in understanding the bacterial response to antibiotic combinations and devising new treatment strategies.

## 1 Introduction

Our society is facing a major crisis with the rise of antimicrobial resistance. As the pace of the discovery of new antimicrobial molecules has slowed down, it is crucial to use current molecules to their full potential. One strategy to maximise the efficiency of antibiotic treatment and minimise the evolution of resistance is to use antibiotic combinations (Chait et al., 2007; Tyers and Wright, 2019). However, such combinations can impact bacterial growth differently depending on their interaction: as the antibiotics interact, their effect on bacterial growth can be stronger (synergistic) or weaker (antagonistic) than expected from the additive effect of each single drug if they act independently (Mitosch and Bollenbach, 2014; Roemhild et al., 2022). Antibiotic interactions are determined quantitatively by comparison to this expected additive effect which can be calculated using two models, Bliss independence (Bliss, 1939) and Loewe additivity (Loewe, 1928, 1953), depending on how the data is collected. Although antibiotic interaction networks have been systematically determined (Yeh et al., 2006; Chevereau and Bollenbach, 2015), the mechanism underlying many interactions have yet to be found or fully quantified.

The interaction between DNA-targeting antibiotics (such as fluoroquinolones) and some translation inhibitors, such as chloramphenicol or tetracycline, has been analysed in detail at the population level. This interaction is antagonistic (i.e. effect weaker than expected) and, in some concentration regimes, can even be suppressive, i.e. the drug interaction results in a higher population growth rate than that observed for cells exposed to one of the drugs alone (Bollenbach et al., 2009). The current understanding of this suppressive interaction is based on combining the known bacterial response to each antibiotic. Fluoroquinolones (such as ciprofloxacin) are bactericidal antibiotics that cause cell death through the inhibition of the activity of the type II topoisomerases. This leads to the accumulation of DNA doublestrand breaks (DSBs) and interferes with chromosome replication and transcription (Bush et al., 2020). In *Escherichia coli* the DNA damage response, called the SOS response, is induced and leads to the expression of approximately 40 genes important for DNA repair, DNA replication and survival (Kreuzer, 2013; Mo et al., 2016). In contrast, translation inhibitors such as tetracycline are bacteriostatic and lead to reduced growth by inhibiting ribosome activities and therefore protein production (Chopra and Roberts, 2001; Poehlsgaard and Douthwaite, 2005; Barrenechea et al., 2021). It has been proposed that under fluoroquinolone treatment, the cells do not appropriately downregulate ribosomal expression resulting in a skewed protein-DNA ratio due to excess and metabolically costly protein production (Bollenbach et al., 2009). When combined, the translation inhibitor partially restores the imbalance in the protein-DNA ratio leading to improved population growth (Bollenbach et al., 2009).

Understanding how drug combinations are affected by changing nutrient environments is important for treating bacterial infections. Indeed, bacterial growth rates are known to vary depending on where in the body the infection is located. For example, *E. coli* doubling times can range from 3 hours in the intestine to approximately 22 minutes in the urinary tract (Myhrvold et al., 2015; Forsyth et al., 2018). The composition of the cell is known to respond to changes in the nutrient environment according to universal constraints on their proteome (Scott et al., 2010; Hui et al., 2015). This can affect the bacterial response to ribosome targeting antibiotics: bacterial susceptibility to tetracycline is growth rate dependent (Greulich et al., 2015) with fast-growing bacteria more affected than slow-growing ones. In addition to changes in cell proteome composition, changing growth rates by varying nutrient composition can impact DNA replication and potentially the efficiency of DNA repair. Fast-growing cells (less than 60 min doubling time) undergo multi-fork DNA replication (Cooper and Helmstetter, 1968) initiating multiple overlapping rounds of replication. This results in higher DNA content per cell, a higher number of replication forks and multiple partial chromosomal copies in fast-growing cells. Because of their higher DNA content, treatment with ciprofloxacin may lead to more DSBs in fast-growing cells, but these cells also have a higher number of potential copies for homologous repair, while slower-growing cells may be limited in their capacity to repair due to a lack of homologous copies of the chromosome. This may lead to growth dependence of the susceptibility to ciprofloxacin, which has indeed been shown at the population level (Smirnova and Oktyabrsky, 2018). Growth dependence of susceptibility to individual antibiotics suggests that antibiotic combinations are also likely affected by changes in nutrient quality, but this has not yet been investigated, especially at the single-cell level.

It is well established that isogenic bacterial cultures show high phenotypic heterogeneity at the single-cell level, including cell-to-cell variability in gene expression, growth rates, stress responses, and susceptibility to antibiotics (Jones and Uphoff, 2021; Vincent and Uphoff, 2021; Sampaio et al., 2022; Brandis et al., 2023; Choudhary et al., 2023). Antibiotics can induce further heterogeneity in the population. For example, we showed previously that DNA damage leads to high heterogeneity in division rates due to variability in SOS expression among single cells and that this can have an impact on the population dynamics (Jaramillo-Riveri et al., 2022). Furthermore, single-cell elongation rates and division rates can become uncoupled under DNA stress in cells that filament, leading to inaccurate optical density readings (Stevenson et al., 2016) and growth rate measurements (Jaramillo-Riveri et al., 2022). Therefore, acquiring single-cell time-resolved data is vital for understanding the dynamic response of bacteria to antibiotic combinations.

Traditionally, population growth rates are determined using bulk-level methods (e.g. optical density or colony-forming unit counting), which measure net growth and population-averaged behaviour. These measurements are a function of cell mass accumulation (which correlates with elongation for rod-shaped bacteria), cell birth/division, and cell death. For the antagonistic drug combination between ciprofloxacin and tetracycline, it is unclear what effect the drug combination has on each of these parameters and which one results in the observed increase in population growth rate. For instance, faster growth than expected under the drug combination could indicate an increase in single-cell elongation, division, survival, or a combination thereof. In this study, we used a microfluidic device called the mother machine to measure each of these parameters separately and determine how the drug combination affects bacteria in three growth conditions. We showed that the antagonism between ciprofloxacin and tetracycline is due to increased survival of cells under the drug combination compared to ciprofloxacin alone and is not due to increased division or mass accumulation. This effect is growth-dependent, with better suppression in fast growth than in slow growth. Quantification of the DNA damage response revealed two sub-populations among the cells dying upon ciprofloxacin treatment, with some cells reaching a very high level of SOS while others had a lower level of SOS similar to cells that survived. The low-SOS cells were more frequent in fast growth conditions and showed increased survival in the antibiotic combination contrary to the high-SOS ones, thus explaining the growth dependence of the antagonistic effect of tetracycline on ciprofloxacin.

## 2 Materials and Methods

### 2.1 Culture conditions

For all microscopy and batch experiments, cell cultures were grown in M9-based media. The composition of the M9 salts was as follows: 49 mM Na_2_HPO_4_, 22 mM KH_2_PO_4_, 8.6 mM NaCl, 19 mM NH_4_Cl, 2 mM MgSO_4_, and 0.1 mM CaCl_2_ (adjusted to pH=7). This was supplemented with either 0.5% w/v glycerol (Sigma-Aldrich, G9012), giving rise to the “gly” medium; 0.5% w/w glucose (D(+)-glucose anhydrous, Scientific Laboratory Supplies, CHE1806), “glu” medium; in the third medium, “glu-aa”, a mix of amino acids (1× MEM Non-Essential Amino Acids and 1× MEM Essential Amino Acids, both manufactured by Gibco™) was added with the glucose. For strains and plasmid construction, cells were grown in LB, or LB agar supplemented with the corresponding selection marker (kanamycin 50 *µ*g/ml). All cultures were grown at 37°C (150 rpm agitation) in 50 ml falcon tubes with no more than 5 ml of liquid volume or 500 ml Erlenmeyer flasks with no more than 50 ml of liquid volume, unless otherwise stated.

Antibiotics used in this study are listed as follows: ciprofloxacin hydrochloride (APExBIO, C5539) and tetracycline hydrochloride (Fisher BioReagents, BP912-100). Antibiotic stocks were made up in sterile distilled water and filter-sterilised (0.22 *µ*m). Stock concentrations were 5 mg/mL for ciprofloxacin (pH-adjusted to 4.62) and 10 mg/mL for tetracycline. Antibiotic stocks were aliquoted into small volumes (*<* 200 *µ*L) and frozen at -20°C. For each experiment, new antibiotic stocks were thawed on the same day and never re-used. Working stocks were diluted using sterile M9-based medium. Fresh tetracycline stocks were made up annually using a new powder batch due to its low long-term stability. When used, antibiotics were always shielded from light.

### 2.2 Strain construction

*Escherichia coli* K-12 MG1655 was used as WT strain in this study. Gene expression reporters (SOS expression: P*_sulA_-mGFP* ; constitutive expression: P*_tet_*_01_*-mKate2* ) were inserted into the genome by clone-integration as described previously (Jaramillo-Riveri et al., 2022). For the *motA* gene deletion strain, a single gene deletion knockout (from the Keio deletion collection (Baba et al., 2006)) was introduced in our strain using P1 transduction. The *motA* gene was removed to inhibit flagellum activity to improve cell retention in the microfluidic microchannels. The antibiotic resistance selection markers were removed by transformation of the pE-FLP plasmid (St-Pierre et al., 2013). After construction, inserts were checked by PCR amplification and sequencing of the modified chromosomal region. The strains, plasmids, and primers used are listed in Supplementary Table S2, S3, and S4, respectively.

### 2.3 Fluorescence microscopy

All images were captured using a Nikon Ti-E inverted microscope equipped with EMCCD Camera (iXion Ultra 897, Andor), a SpectraX Line engine (Lumencor) and a 100× Nikon TIRF objective (NA 1.49, oil immersion). Nikon Perfect-Focus system was used for continuous maintenance of focus. The filter set for imaging mGFP consisted of ET480/40× (excitation), T510LPXR (dichroic), and ET535/50m (emission); whereas for mKate2 the set ET572/35× (excitation), T590LPXR (dichroic), and ET632/60m (emission) was used. Filters used were purchased from Chroma. mGFP fluorescence was measured using 80 ms exposure, whereas mKate fluorescence was imaged for 100 ms, both at minimal camera gain and maximum lamp intensity. Brightfield images were also acquired at each frame using a 20 ms exposure. The microscope and temperature chamber was turned on to 37°C at least 2 hours before imaging. The microscope was controlled from MATLAB via MicroManager (Edelstein et al., 2014) using a custom-made user interface developed by Sebastian Jaramillo-Riveri (Jaramillo-Riveri, 2019). The code is accessible at https://gitlab.com/MEKlab/MicroscopeControl. Images were saved in *tiff* format as one file per fluorescence channel per frame.

### 2.4 Mother machine experiments

#### 2.4.1 Microfluidic device design and fabrication

The design and fabrication of the mother machine device was described previously (Jaramillo-Riveri et al., 2022). The silicone wafers were manufactured by Micro Resist Technology GmbH. Microchannel dimensions were optimised for each medium and are indicated in Supplementary Table S5.

#### 2.4.2 Culture preparation and experimental design

The culture was prepared as follows: Cells were inoculated in 5ml LB from a -80°C stock and grown overnight. The next morning (day of experiment), cells were sub-cultured (1:1000 dilution factor) into 2X 10 mL M9-based medium in 50 mL falcon tubes and incubated at 37°C with shaking (150 rpm). When the culture reached an OD_600_ of 0.2, cells were harvested for inoculation in the mother machine. For the slow growth medium (gly), an extra step was added: After 1:1000 dilution from the LB overnight into M9-based medium (late afternoon), the culture was incubated overnight. The next morning, cells were diluted again (1:200) and allowed to grow until OD_600_ = 0.2.

Once the culture was ready for inoculation, Tween-20 Surfact-Amps detergent solution (Thermo Scientific, 85113) was added to prevent cell clumping (0.01% final concentration) before being concentrated 100-fold by centrifugation (4000 rpm for 5 min). Before sample loading, the chip was passivated with Tween-20 (0.01% final concentration) for at least 1 hour. The concentrated cell culture was briefly vortexed and then injected into the inlets of the microfluidic chip using a 1 mL syringe with a 21-gauge blunt needle (OctoInkjet), while taking care not to damage channel inlets. The channel inlets were sealed temporarily using adhesive tape and left at 37°C for approximately 30 min to allow cells to diffuse into the microchannels. To further assist cell loading, the cells were then spun into the microchannels by centrifugation at 4000 rpm (3220 g) for 5 min using a custom-built mount. Centrifugation was essential for loading of non-motile cells. The chip was then mounted on the microscope using a custom-built mount and the loading efficiency was evaluated. Before and after each experiment, the inlet and outlet tubing (0.44 mm ID) were sterilised with 1% bleach, followed by 10% ethanol, rinsed with Milli-Q water, and then primed with fresh medium. Tween-20 (0.01% final concentration) was also added to the medium reservoir to prevent cell adhesion and biofilm formation within the tubing and device. Inlet and outlet tubing was then connected to the inlet and outlet channels of the microluidic device via 21-gauge blunt needles with the medium reservoir on one end and effluent reservoir on the other end. Tubing was connected to the reservoirs via a custom-built manifold. Inlet tubing was also connected to three-way valves (two inlets, one outlet) which were used to switch between no-drug and antibiotic influent reservoirs. Valves were controlled automatically via MATLAB communication with a data acquisition device (DAQ, Measurement Computing, USB-1408FS). Valve-outlet tubing was then connected to a peristaltic pump (Ismatec IPC ISM932D) which ‘pushed’ (positive-pressure flow is advantageous as it avoids feature collapse due to negative-pressure driven flow) medium through the microfluidic chip into the effluent flask.

Once cells were loaded into the microfluidic device, medium flow was used to flush cells out the main trench using a peristaltic pump at a flow rate of 1.5-2 mL/h and then lowered to 1 mL/h for the duration of the experiment. All flow channels received fresh medium without antibiotic for at least two hours to allow for recovery. Images were then acquired at 5 min intervals for glu-aa and glu media, and 10 min for gly medium. When imaging started, all channels received medium without antibiotic for a further 6 hours, followed by 12 hours of antibiotic exposure, and then switched back to medium without antibiotic for at least 10 hours, unless otherwise stated. The pre-antibiotic period ensures all cells have acclimated to the system and are undergoing balanced exponential growth. The post-antibiotic period (or recovery period) allows the resuscitation of persister cells, if present. Antibiotics solutions were prepared on the day of the experiment to the desired concentration in M9-based medium in separate 50 mL falcon tubes. These were kept at 37°C for the duration of the experiment and wrapped in foil to protect from light. The microfluidic device has four independent flow channels. Thus, for each experiment, four treatments ran simultaneously: A no-drug control, CIP 3 ng/mL, TET 0.2 *µ*g/mL, and the drug combination (CIPTET) at the same concentrations. At least two experimental replicates were performed on different days, except for CIP in glu-aa, the control in gly, and TET in gly, where three replicates were performed.

#### 2.4.3 Image analysis

Cell segmentation and lineage tracking were performed using BACMMAN run in Fiji (Ollion et al., 2019). The BACMMAN configuration was adapted to segment and track cells based on fluorescence images. Images (*tiff* format) were imported in BACMMAN and pre-processed prior to segmentation. This included image rotation to ensure channels were vertically oriented and the channel opening was at the bottom, and cropping of images to include only the area consisting of microchannels. Background subtraction was performed for both fluorescent channels using ImageJ’s subtract background algorithm (rolling ball method, radius=8). Joint segmentation and tracking of cells were performed using a DistNet model trained on our datasets. DistNet is based on a deep neural network (Ollion and Ollion, 2020). Segmentation was performed on the mKate fluorescence channel. Segmentation parameters were optimised for each dataset. Curation of segmentation and tracking was carried out using BACMMAN’s interactive graphical interface. Although automated segmentation and tracking was mostly accurate, occasionally errors were produced. Thus, almost every lineage was manually checked, with CIP exposure datasets requiring the most curation mainly due to excessive elongation of cells. At this point obvious irregularities were removed including mother cells that did not grow for the duration of the experiment and those that were already excessively elongated at the beginning of the experiment. Positions with channel deformities and where loss of focus occurred were also discarded.

Single-cell fluorescent intensities were calculated by dividing the sum of pixel intensity values by the cell area. Cell length was determined as the maximal distance between two points of the cell contour (FeretMax). Elongation rates were calculated per cell-cycle (generation) by fitting an exponential function, *S_t_* = *S*_0_*e^µt^*, on the estimated length of cells, where *µ* is the elongation rate, *S*_0_ is the length at birth, *S_t_* is the length at time *t*. A minimum of 3 frames per generation was imposed as a fitting constraint. Division rates were calculated as the inverse of the interdivision time. The latter was determined as the time interval between two divisions. Cell fluorescence and morphology measurements were then exported from BACMMAN into csv files for further analysis.

#### 2.4.4 Data analysis

Data analysis and plotting was conducted using Python in Jupyter notebooks. The first 4 hours of all experiments were discarded to ensure only cells at steady-state were analysed. Thus, each dataset begins with 2 hours of no-drug medium before antibiotic exposure. For antibiotic treatment experiments, we removed lineages that were initially filamented or non-growing in the first 2 hours. Some cells that filamented extended beyond the length of the microchannels (classified as hyper-filamented cells). We only tracked a cell that filamented up until it reached the maximum microchannel length since cell length measurements beyond that are invalid. Classification of cell fate and sub-classification of low & high-SOS inducers is described in the Supplementary Methods. Survival analysis was carried out using a Kaplan-Meier estimator (Kaplan and Meier, 1958) described in detail in the Supplementary Methods. The Bliss independence expected survival fraction (*Sf* ) presented in Figures 3D and 5G-H was derived from the log-transformed pairwise sums of the replicate *Sf* s in each group: *ln*(*Sf_CIPij_*) + *ln*(*Sf_TETij_* ), where *i*=1,2,3 for the three growth media and *j*=1,…,*n_i_*, where *n_i_* is the number of replicates in the *i*th group. The level of suppression calculated in Figure 5I was defined as Σ = (*S̄f_CIP_* _-*TET*_ *− S̄f_CIP_*)*/S̄f_CIP_*, where *S̄f_CIP_* _-*TET*_ is the mean survival fraction under the drug combination and *S̄f_CIP_* is the mean survival fraction under CIP which is antagonised by TET. Equation adapted from Bollenbach et al. (2009).

### 2.5 Statistical analysis

Statistical analysis of the Kaplan-Meier survival curves were conducted using the Log-rank test. For the survival fractions at the end of the experiment, a two-sided *t* -test was used to test for significant differences in means. To test the null hypothesis of drug independence according to Bliss, we used an ANOVA model described by Demidenko and Miller (2019). This was used to determine if the observed *Sf* under CIP-TET was significantly higher than the expected *Sf* according to Bliss independence. All statistical analysis was performed in Python.

## 3 Results

### 3.1 The single-cell response to sub-lethal concentrations of ciprofloxacin and tetracycline is highly heterogeneous

To determine the conditions where the combination of ciprofloxacin (CIP) and tetracycline (TET) is antagonistic (or suppressive), we performed an initial characterisation in bulk culture using a defined medium that supports fast growth (M9 based medium supplemented with glucose and amino acid, here referred to as glu-aa). Analysis of growth rate using OD_600_ measurements confirmed the suppressive interaction of TET and CIP in a concentration range of 4.5-13.5 ng/ml for CIP and 0.2-0.6 *µ*g/mL for TET (Supplementary Figure S1A, B). However, initial tests in a microfluidic device called the ‘mother machine’, which allows the monitoring of hundreds of single-cell lineages under defined growth conditions (Wang et al., 2010), revealed that when exposed to 5 ng/ml CIP, cells undergo high levels of filamentation. Filamentation under CIP treatment is expected since CIP creates DNA damage and leads to the induction of cell division inhibitors as part of the SOS response (Kreuzer, 2013). However, the observed level of filamentation resulted in a too large number of cell lineages being lost. We, therefore, used 3 ng/ml CIP and 0.2 *µ*g/mL TET for all the subsequent experiments: this concentration allows the detection of the antagonistic relationship between the two antibiotics whilst limiting cell filamentation (Supplementary Figure S1C), which facilitates imaging of individual cells in the microfluidic device.

We first sought to evaluate the single-cell physiological response of *E. coli* to exposure to CIP and TET separately at the concentration previously determined. We cultivated cells in the mother machine and after adaptation measured the bacterial growth for 2 hours. We then added the relevant antibiotic for 12 hours, removed it and monitored the cells for a further 10 hours to allow recovery from antibiotic treatment (Figure 1A). The bacteria were imaged by fluorescence microscopy and segmented and tracked automatically (see Materials and Methods). We only tracked the lineage of the cell at the end of the microfluidic microchannel (the ‘mother’ cell). We measured the single-cell elongation rates, division rates, cell length, and gene expression using a constitutively expressed gene integrated on the chromosome (*P_tetO_*_1_-*mKate*2). Additionally, the strain carried a transcriptional reporter for the SOS response (*P_sulA_*-*mGFP* ) (Jaramillo-Riveri et al., 2022). As shown in example kymographs (Figure 1B), cells in the no-drug condition grew and divided for the entire duration of the experiment. When exposed to TET, the cells became slightly smaller, but continued to elongate and divide (Figure 1C), as expected when cells are exposed to sublethal concentrations of a bacteriostatic antibiotic. In contrast, when the bacteria were exposed to CIP, we detected a high level of heterogeneity in their response. Some cells continued growth and division, while others filamented to varying degrees before either resuming division (Figure 1D) or stopping growth and division for the remainder of the experiment (Figure 1E). Additionally, some cells ceased growth and division without filamenting.

**Figure 1:**
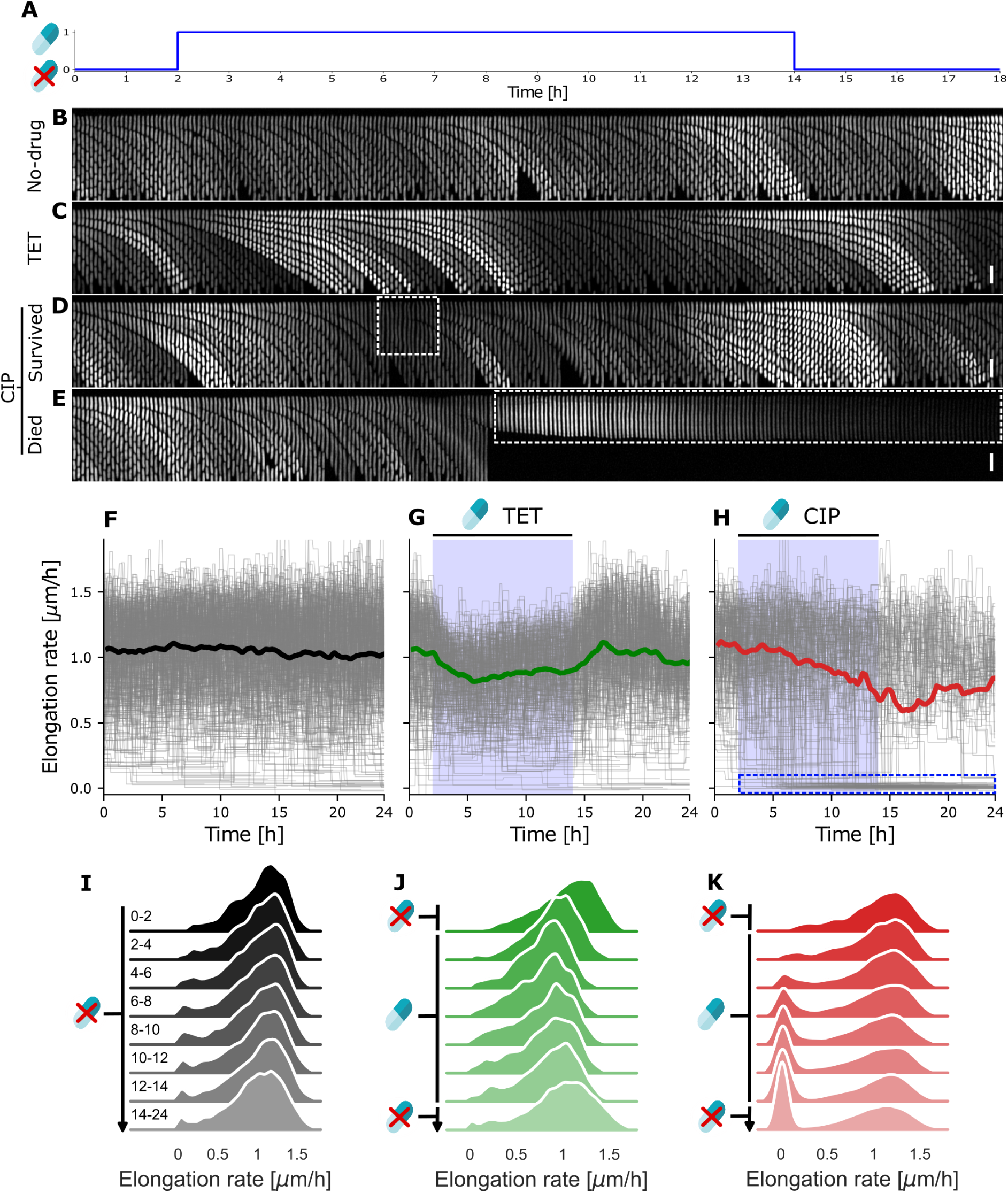
Single-cell response under sub-lethal antibiotic treatment is heterogeneous. **A**. Schematic indicating when antibiotic was introduced (hour 2) and then removed (hour 14). **A-E**. Example kymographs showing growth under no-drug, 0.2 *µ*g/mL tetracycline and 3 ng/mL ciprofloxacin treatment. Shown is fluorescent intensity from the *P_tetO_*_1_-*mKate*2 constitutive reporter. Under CIP treatment, some cells continued growth and division with some filamentation (D, white box), while others stopped division (E, white box). Kymographs were constructed as a montage of frames with 5-minute intervals from a single microchannel. Scale bar represents 5 *µ*m. **F-H**. Single-cell trajectories of elongation rates under no-drug (F, n = 334 cells), 0.2 *µ*g/mL tetracycline (G, n = 237), and 3 ng/mL ciprofloxacin (H, n = 191) treatment. Antibiotics were introduced between hours 2-14, indicated by the blue shaded area. Grey lines represent individual mother cell lineage trajectories. Solid lines represent the median of the population. Shown is data from one experimental replicate for growth in glu-aa medium. **I-K**. Distributions of single-cell elongation rates for the no-drug control (I), TET treatment (J) and CIP treatment (K) drawn from sequential two-hour periods under different antibiotic treatments in the glu-aa growth medium. Distributions are kernel density estimates of the underlying histogram pooled from at least two experimental replicates. Time periods, in hours, are indicated to the left of each distribution. Antibiotics were introduced from hours 2-14 as indicated.

We then quantified single-cell elongation rate for all cells under CIP, TET, and the no-drug control. Elongation rate trajectories under drug-free conditions remained stable over the duration of the experiment (Figure 1F). Under TET treatment, elongation rates initially decreased before levelling off to a median level approximately 16% below pre-antibiotic treatment. When TET was removed, elongation rates recovered to pre-antibiotic levels (Figure 1G). To compare the elongation rates over time, we plotted their distributions over 2-hour windows for the duration of the experiment. The response appeared homogeneous across the population in the no-drug and TET conditions, suggesting that there was limited heterogeneity under these conditions (Figure 1I, J). Under these conditions, we detected a very small population that ceased growth toward the end of the experiment, suggesting age-related death as has been reported before (Wang et al., 2010; Yang et al., 2019).

In contrast, elongation rates under CIP treatment were highly heterogeneous (Figure 1H). A large fraction of the cells did not show any change in elongation rates during CIP exposure. However, a second population of cells stopped elongation entirely (trajectories near 0 *µ*m/h; Figure 1H blue box). We did not observe any significant recovery of cells’ elongation within the 10 hours of drug-free growth post antibiotic treatment, suggesting that these cells had died. This second population appeared clearly in the distributions shown in Figure 1K from the 4-6 hour window (i.e. 2 to 4 hours after antibiotic exposure), and its proportion increased over time. To further check whether these cells were indeed dying, we measured the expression of the constitutively expressed mKate2 marker: all showed an exponential decrease in gene expression after elongation stopped, further indicating that protein synthesis had stopped and that these cells were not metabolically active (Supplementary Figure S5-S7F). We also checked whether a longer recovery period of 36 hours would lead to some cells resuming growth, and found no significant recovery. Taken together, these results indicate that cells that have abruptly stopped elongation are ‘dead’, similar to the definition used in previous studies (Robert et al., 2018; Vincent and Uphoff, 2021). Therefore, we conclude that even at the very low concentration of 3 ng/ml CIP (whereas the MIC is typically 12 ng/ml (Bollenbach et al., 2009; Pribis et al., 2019)) we observe a significant amount of cell death. This is not surprising given that CIP is bactericidal and has been previously observed at the population level (Coates et al., 2018).

### 3.2 The CIP-TET combination is suppressive due to improved survival of bacterial cells

We then exposed cells to a combination of CIP and TET at the same concentrations as before. Single-cell elongation rates under the drug combination uniformly decreased, similarly to cells treated with TET alone (Figure 2A, B). Median elongation rates of the growing population at the end of drug treatment decreased by approximately 14% (versus 16% in TET alone). However, in contrast to cells exposed to CIP, we observed a marked decrease in the proportion of cells that stopped growth and division under the drug combination (Figure 2C). Moreover, we did not observe an increase in elongation rates (when compared to TET exposure alone) for the growing population when TET was added to CIP (Figure 2C). This indicates that, at the single-cell level, there is no suppressive effect on elongation rates under the drug combination, but that the drug combination acts by decreasing the number of cells that die. Therefore, the increase in population growth rate that we observed in bulk experiments (Supplementary Figure S1) and has been reported before (Bollenbach et al., 2009), is not due to better growth (i.e., increase in mass/volume of cells) in CIP-TET but due to lower cell death under the antibiotic combination.

**Figure 2:**
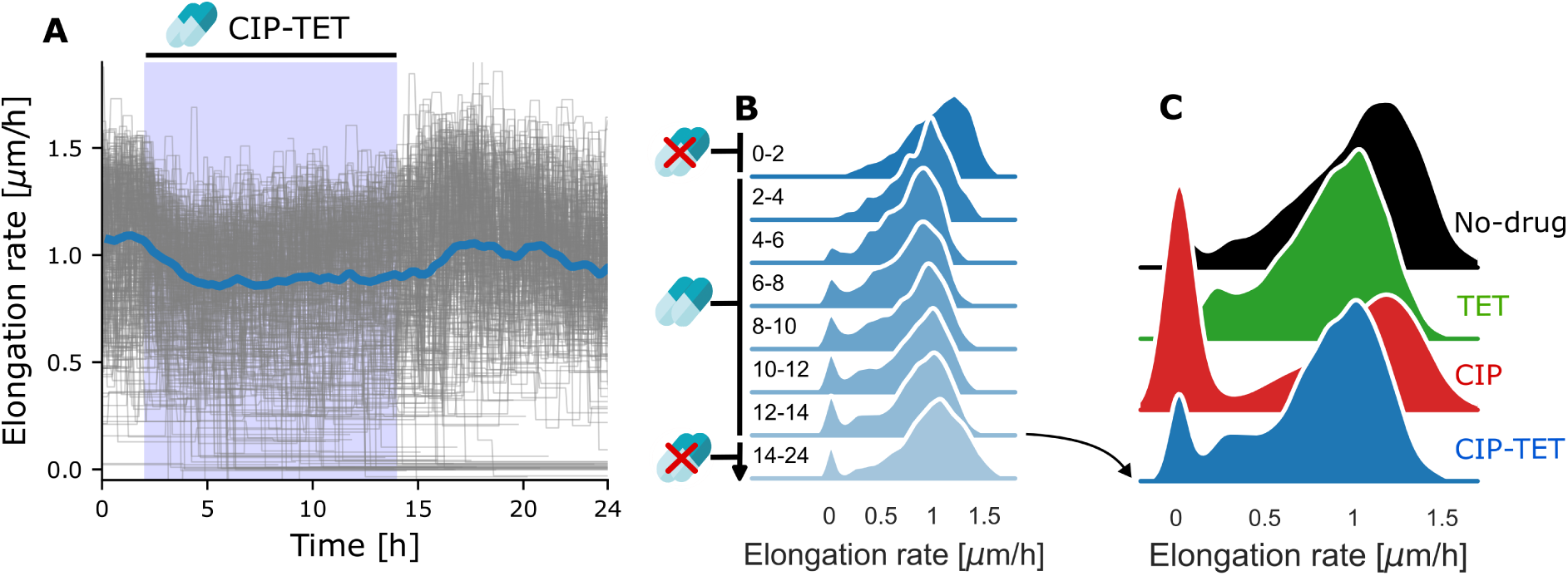
The drug combination improves cell survival with no increase in single-cell elongation rates. **A**. Single-cell trajectories of elongation rates under the combination of 0.2 *µ*g/mL tetracycline and 3 ng/mL ciprofloxacin treatment (n = 304 cells). Antibiotics are introduced between hours 2-14, indicated by the blue shaded area. Grey lines represent individual mother cell lineage trajectories. Solid lines represent the median of the population. Shown is data from one experimental replicate for growth in glu-aa medium. **B**. Distributions of single-cell elongation rates drawn from sequential two-hour periods under CIP-TET treatment in the glu-aa growth medium. Distributions are kernel density estimates of the underlying histogram pooled from at least two experimental replicates. Time periods, in hours, are indicated to the left of each distribution. Antibiotics were introduced from hours 2-14 as indicated. **C**. Distributions of single-cell elongation rates under different treatment conditions during the final 2 hours of antibiotic treatment. As in C, distributions are kernel density estimates of the underlying histogram pooled from at least two experimental replicates.

**Figure 3:**
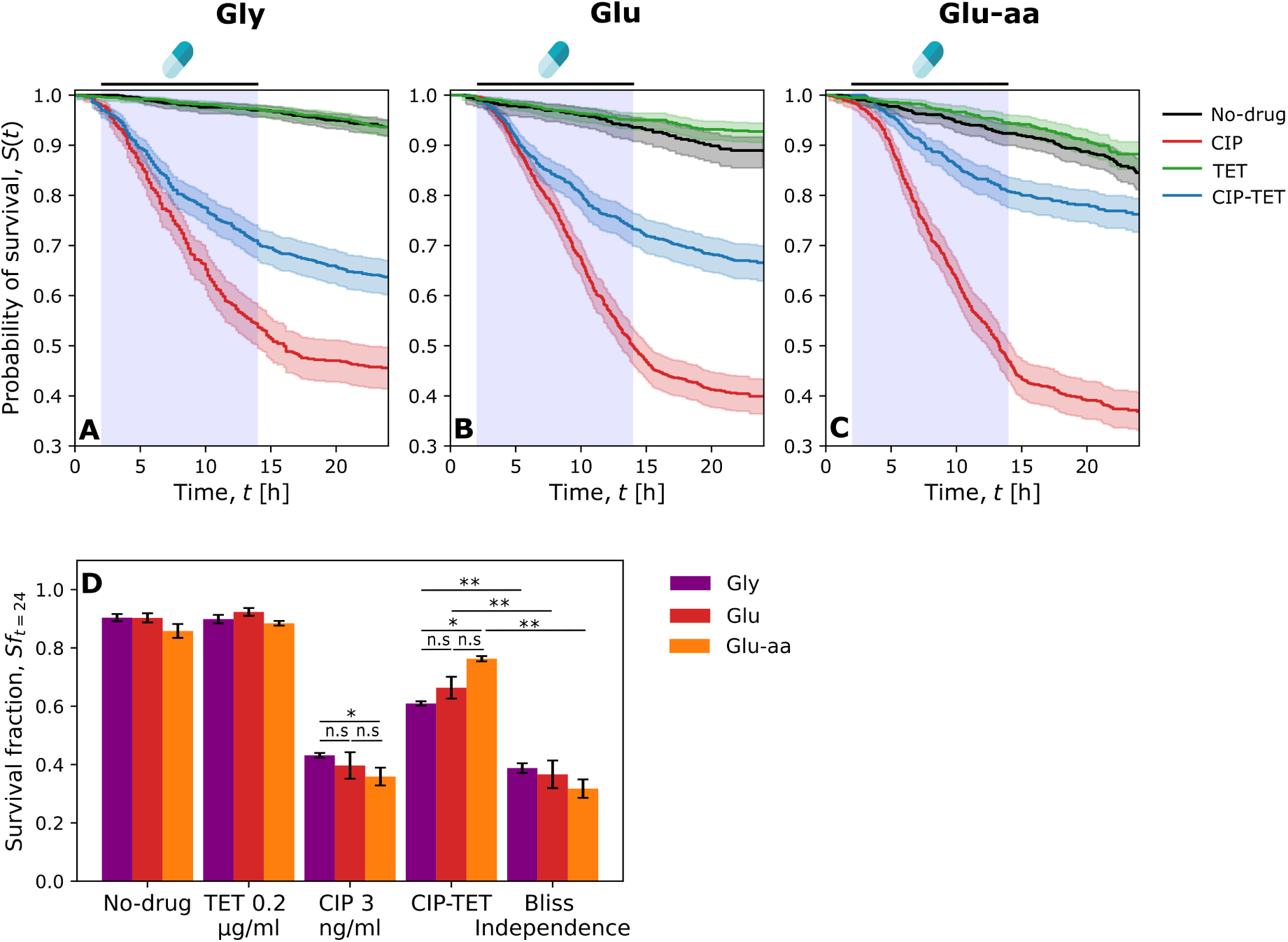
The CIP-TET combination improves survival in all growth conditions. **A-C**. Survival probability curves under different antibiotic treatments in three different growth media. Survival probabilities (*S_t_*) were calculated using the Kaplan-Meier estimator and mother cell survival times pooled from at least two experimental replicates consisting of 420-1005 mother cell lineages. Antibiotics were introduced between hours 2-14 (blue shaded area). Shaded area represents 95% confidence intervals of the estimated survival probability. Survival curves under CIP and CIP-TET are significantly different in all growth media (Log-rank test, *p<*0.005). Survival curves for no-drug control and TET are not significantly different. **D**. Survival fractions (*Sf* ) for low-SOS cell death (G) and high-SOS cell death (H) were quantified at the end of the experiment under antibiotic mono-exposure and drug combination in three different growth media. See Materials and Methods for calculation of Bliss independence and statistical analysis. Significant difference of means among and between the CIP and CIP-TET groups were calculated using a two-sided *t* -test. Significant differences are indicated: \*\**p<*0.005, \**p<*0.05, n.s.= not significant. All survival fractions under CIP-TET were significantly higher than under CIP alone (not shown on figure). Error bars represent the standard error of the mean.

### 3.3 The CIP-TET combination is suppressive in all growth conditions

To check whether survival was affected by growth rate, we compared the survival of bacteria under CIP, TET, and CIP-TET exposure in three different growth conditions using the same concentration as before. In addition to the glu-aa condition, we used an M9-based medium supplemented with glucose (referred to as ‘glu’) or glycerol (referred to as ‘gly’). As expected, the median elongation rate was higher in glu-aa than in glu and gly (1.13, 0.54, & 0.28 *µ*m/h median elongation rates, respectively) in the no-drug control.

To quantify the survival of individual cells more precisely, we devised a simple algorithm to classify a lineage as having survived or died (see Supplementary Methods). Following the classification of lineage fate, we generated survival curves (Figure 3A-C) using the Kaplan-Meier (K-M) estimator (Kaplan and Meier, 1958). As expected, most cells survived in the no-drug control and in the TET treatment. Similar to our initial observation in glu-aa, we observed a small amount of death in glu and in gly at the end of the experiment, likely due to ageing (Wang et al., 2010; Yang et al., 2019). In contrast, under CIP treatment, we observed reduced survival in all growth conditions: most cells died during antibiotic treatment, or within a few hours after antibiotic removal, probably due to residual antibiotic or residual DNA damage in the cells. After the antibiotic was removed, the death rate decreased. Importantly, CIP was more effective in glu-aa (fast growth condition) than in gly, in keeping with what has been observed previously at the population level (Smirnova and Oktyabrsky, 2018). Under the antibiotic combination, single-cell survival significantly improved (Log-rank test, *p<*0.005) in all growth conditions (Figure 3A-C, compare red and blue curves). When measuring mean survival fractions at the end of the experiment (Figure 3D), cells under CIP showed 43%, 38% and 36% survival for gly, glu & glu-aa, respectively. Under the CIP-TET combination, the proportion of cells that had survived was 61%, 66% and 76% for gly, glu & glu-aa, respectively, which is significantly higher in all growth conditions (two-sided *t* -test, *p<*0.05). We calculated the expected effect (*Survival_CIP_ Survival_T ET_* = *Survival_CIP_ × Survival_T ET_* ) of the two antibiotics under the Bliss independence hypothesis (Bliss, 1939; Demidenko and Miller, 2019) (Materials and Methods). In all growth conditions, we observed that *Survival_CIP_* _-_*_T ET_ ≫ Survival_CIP_ Survival_T ET_* which indicates that the interaction is antagonistic on cell survival. Moreover, as survival was higher than in CIP treatment alone, we conclude that the drug interaction is suppressive (Roemhild et al., 2022) in all growth conditions, even at the very low CIP concentration that we used.

### 3.4 The SOS response is highly heterogeneous

In order to better understand the underlying mechanism that leads to cell death, we measured the SOS response of cells exposed to the different antibiotic treatments using the *P_sulA_*-*mGFP* transcriptional reporter (Figure 4). We calculated the average fluorescence per cell area (which we refer to as “SOS expression”) as a proxy for GFP concentration for each cell at each time point (Jaramillo-Riveri et al., 2022). To compare SOS expression in all conditions, we analysed the SOS expression distribution over the final two-hour window of antibiotic exposure, during which the SOS expression had reached a steady state. In the no-drug control and under TET exposure, the cells did not induce SOS significantly in any of the growth conditions. However, as expected, under CIP, there was an increase in SOS expression (median SOS expression for no-drug control: 118, 95, 93 a.u. in glu-aa, glu, & gly; median SOS expression for CIP: 227, 473, 462 a.u.). Moreover, we noticed that the SOS response was highly heterogeneous, with a sub-population of cells reaching an SOS level approximately 10 times higher than the rest of the population (Figure 4B, C, red lines) which we will refer to as the high-SOS population below (we refer to the other population as the “low-SOS” population). The high-SOS population was much more abundant in slow growth conditions (glu & gly) and hardly noticeable in glu-aa. In the CIP-TET condition, the high-SOS population, surprisingly, did not seem to be affected (Figure 4B, C). The low-SOS population did not show not much change in SOS expression in the glu-aa medium (Figure 4A). However, in glu and gly we observed a decrease of SOS expression in the low-SOS population as can be expected from the impact of TET on protein production (compare main peak of red and blue lines in Figure 4B, C). This observed growth dependence of SOS expression under CIP-TET is expected: degradation of LexA upon DNA damage leads to SOS promoters to become de-repressed and behave similar to constitutive promoters; constitutive protein expression is more affected in slow growth than in fast growth under TET inhibition because of the well established universal proteome constraints (Scott et al., 2010; Hui et al., 2015).

**Figure 4:**
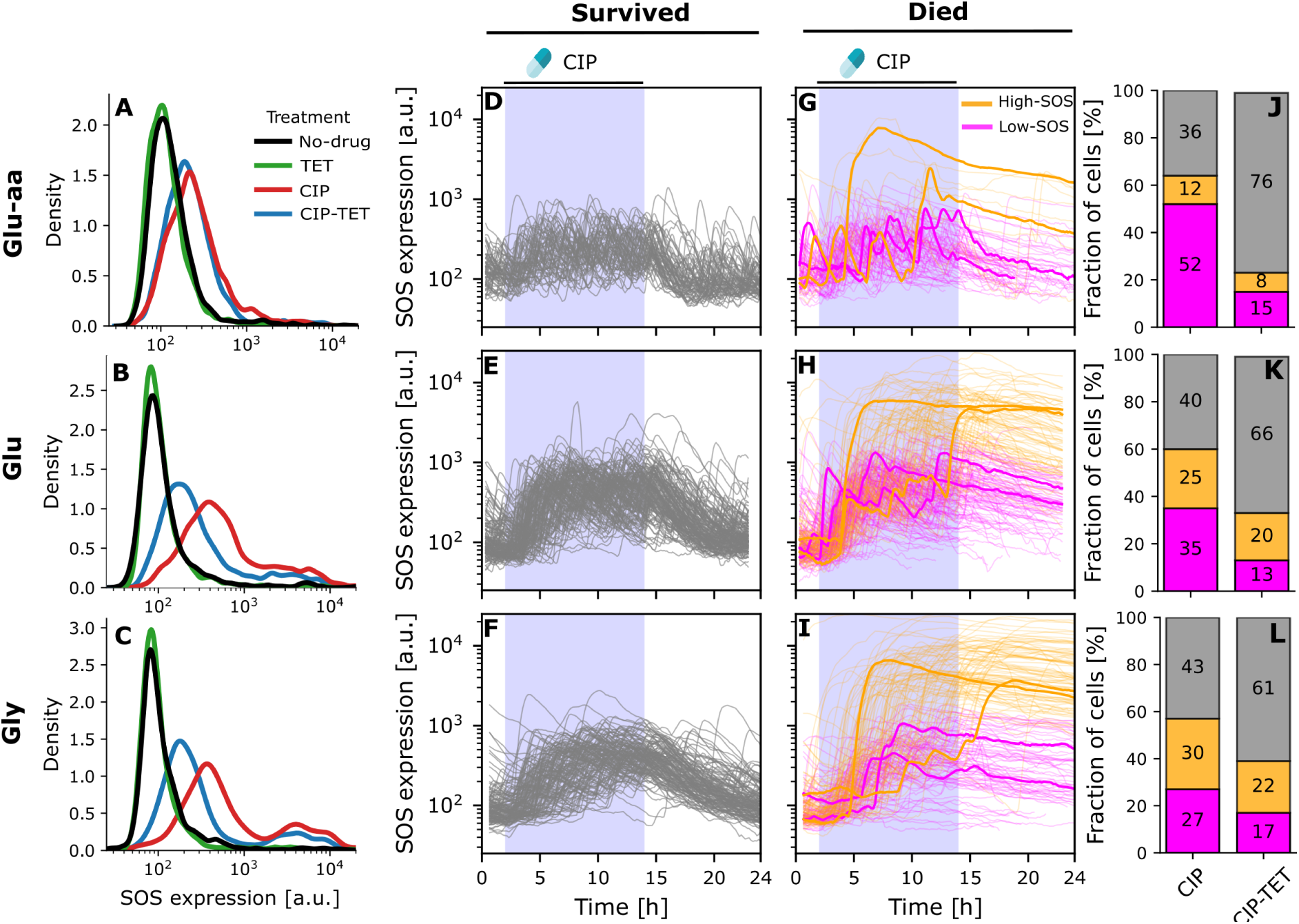
The SOS response is highly heterogeneous under CIP treatment. **A-C**. Distributions of single-cell SOS expression from *P_sulA_*-*mGFP* for the no-drug control (I), TET treatment (J) and CIP treatment (K) drawn from the final 2 hours of antibiotic treatment in different growth conditions. Distributions are kernel density estimates of the underlying histogram pooled from at least two experimental replicates. **D-F**. Single cell trajectories of SOS expression for surviving cells under CIP treatment in different growth conditions. CIP was introduced between hours 2-14 (blue shaded area). **G-I**. Single cell trajectories of SOS expression classified by expression level (low-SOS: magenta and high-SOS: orange) for dead cells under CIP treatment in different growth conditions. Example trajectories are highlighted to illustrate the behaviour of the two sub-populations. Shown is data from one experimental replicate for clarity. **J-L**. Fraction of cells classified as survived (grey), low-SOS dead (magenta), and high-SOS dead (orange) over the entire experiment under CIP and CIP-TET treatment under different growth conditions.

**Figure 5:**
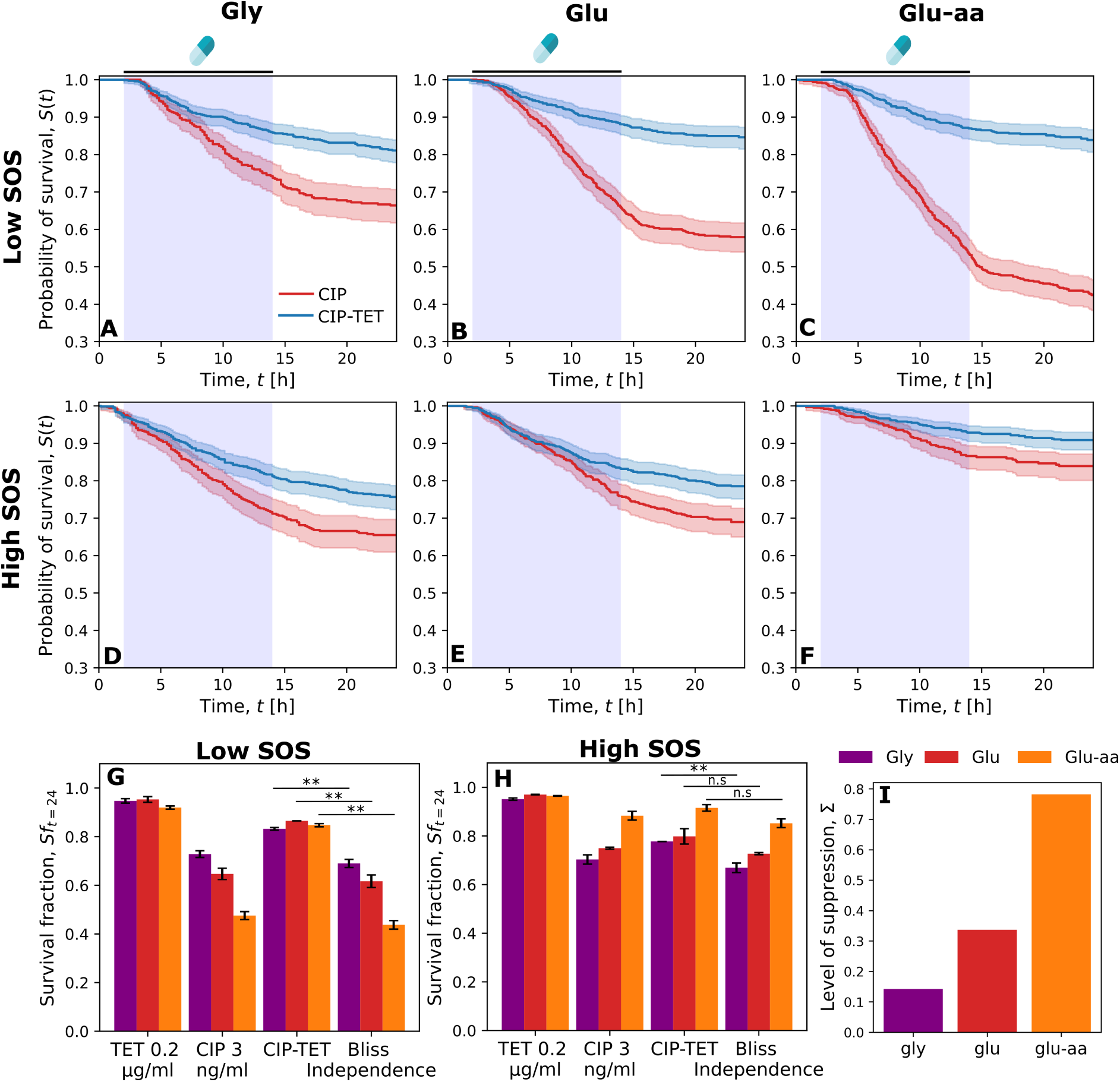
The CIP-TET combination improves the survival of low-SOS cells in a growth-dependent manner. **A-F**. Survival probability curves for low-SOS cells (A-C) and high-SOS cells (D-F) under CIP and CIP-TET treatments in three different growth media. Survival probabilities (*S_t_*) were calculated using the Kaplan-Meier estimator and mother cell survival times pooled from at least two experimental replicates. Antibiotics were introduced between hours 2-14 (blue shaded area). Shaded area represents 95% confidence intervals of the estimated survival probability. Survival curves under CIP and CIP-TET are significantly different in all growth media (Log-rank test, *p<*0.005). **G-H**. Mean survival fractions (*Sf* ) for low-SOS cell death (G) and high-SOS cell death (H) quantified at the end of the experiment under antibiotic mono-exposure and drug combination in three different growth media from at least two experimental replicates. See Materials and Methods for calculation of Bliss independence and statistical analysis. Significant differences are indicated: \*\**p<*0.005, n.s.= not significant. Error bars represent the standard error of the mean. **I**. Level of suppression of low-SOS cell death quantified in three growth conditions (see Materials and Methods for calculation).

To test whether the level of SOS expression could be indicative of the fate of the cells, we evaluated the SOS expression among cells that had survived or died under CIP exposure. Among the survivors, very few cells reached a high-SOS level in any of the growth conditions, as shown in Figure 4D-F. Moreover, their SOS signal returned within a 5-hour window to their pre-antibiotic level upon removal of CIP suggesting that DNA damage had been repaired. The cells that died exhibited a markedly different behaviour: their SOS level peaked, followed by an exponential decrease, consistent with a stop in protein production and exponential decay due to bleaching (similar to what we observed for the constitutively expressed protein mKate). In these cells, we did observe two sub-populations, with some cells reaching a high SOS level (yellow trajectories in 4G-I) whilst others died after expressing a low level of SOS (magenta trajectories). We therefore decided to sub-classify the cells that died into two categories depending on their SOS response level (see Supplementary Methods) (Figure 4G-I, left column). The fraction of high-SOS cells was only 12% in glu-aa but increased to 25% in glu and 30% in gly, suggesting that in these conditions a large population of cells induce SOS very strongly before dying.

We then performed the same analysis for the CIP-TET drug combination: overall, more cells survived as expected from the suppressive interaction between CIP and TET. Among cells that died, we still observed two SOS sub-populations (Supplementary Figure S8). However, the proportion of low-SOS and high-SOS cells was very different from the CIP treatment alone, suggesting these two sub-populations do not react similarly to the CIP-TET combination. Indeed, 52% of the total population died and had a low level of SOS under CIP exposure in glu-aa, but these cells represented only 15% of the population in CIP-TET (Figure 4I). Similarly, in the other growth conditions the proportions of cells that died and were low-SOS was reduced under CIP-TET (Figure 4K-L). This suggests that exposure to the combination of antibiotics increases the survival of the cells that do not reach a high-SOS level (Figure 4J-L, right column magenta). In contrast, the high-SOS sub-population was comparatively much less reduced under the drug combination (Figure 4J-L, yellow), going from 12% to 8% in glu-aa, 25% to 20% in glu and 30% to 22% in gly. Taken together, these results suggest that there are likely two different responses to DNA damage among cells that die: one that reaches moderate levels of SOS expression and is affected by TET treatment and a second one which reaches much higher levels of SOS and is hardly sensitive to TET exposure.

### 3.5 The CIP-TET combination improves the survival of low-SOS cells in a growth-dependent manner

To quantify the importance of the two sub-populations to overall survival, we measured the survival probability of the population in each growth medium separating the contribution of the low and high-SOS cells (Figure 5A-F). A significant improvement in cell survival was observed for both low and high-SOS cells in all growth conditions (Log-rank test, *p<*0.005). Using the survival fractions for low-SOS cells at the end of the experiment, we observed a significant increase under the drug combination compared to the Bliss independence expectation (Figure 5G), indicating a suppressive interaction in all conditions. However, we did not observe a significant improvement in survival for the high-SOS cells, except for the gly medium (Figure 5H). This suggests that the drug combination is not, or only slightly, antagonistic for the high-SOS cells.

We noticed that the relative increase in survival appeared to be higher in faster growth conditions than in slow growth conditions for the low-SOS cells. To compare the level of suppression in each growth condition for these cells, we calculated the proportional increase in survival under CIP-TET compared to CIP only as Σ = (*Sf_CIP_* _-*T*_ *_ET_ − Sf_CIP_* )*/Sf_CIP_* , similarly to what has been proposed by Bollenbach et al. (2009). The level of suppression was higher in glu-aa (fast growth condition) than in gly - an increase in Σ from 0.14 to 0.78 (Figure 5I), suggesting that suppression of cell death is growth rate dependent for the low-SOS cells. Taken together, these results suggest that the low-SOS population is protected by treatment with the combination of antibiotics in a growth-dependent manner.

## 4 Discussion

In this work, we used a “mother machine” to quantify at single-cell level the impact of treatment to a sublethal concentration of ciprofloxacin in combination with tetracycline. We observed that, as expected for a bactericidal antibiotic even at sub-MIC concentration (Coates et al., 2018), treatment with CIP led to significant cell death in a growth-dependent manner. The CIP-TET combination markedly increased cell survival, revealing that the improved growth observed at the level of bacterial populations, which underlies the suppressive interaction of CIP and TET (Bollenbach et al., 2009), is due to improved survival and not improved single-cell growth rates. We also observed that suppression was stronger in faster growth conditions due to a combination of two effects: growth dependent suppression for the low-SOS cells under CIP-TET and a high frequency in slow growth of the high-SOS cells for which antagonism is very weak.

Our single-cell results indicate that it is important to distinguish two different sub-populations amongst the bacteria that die because of CIP treatment. The first one exhibits a very high level of SOS induction (approximately 10 times more than the rest of the population) before dying. It is much more abundant in slow growth (glu, with a doubling time *≈* 82 minutes and gly, with doubling time *≈* 185 minutes) than in fast growth conditions (glu-aa, doubling time *≈* 40 minutes) (Supplementary Table S1). This might be surprising at first glance, as we recently reported that the rate of high SOS induction in response to a replication-dependant break was higher in fast growth (Jaramillo-Riveri et al., 2022). However, when treating bacteria with ciprofloxacin, DSBs can happen anywhere on the chromosome and are not necessarily replication-dependent (Zhao et al., 2006). Therefore, we do not expect a higher level of breaks in fast-growing cells for this type of DNA damage. Moreover, if a DSB occurs in a region of the chromosome that has not been replicated, its repair by homologous recombination would not be possible. In slow growth conditions, where the cell cycle is longer than 60 minutes, cells do not undergo multi-fork replication and a larger proportion of the chromosome is unreplicated during the cell cycle (Cooper and Helmstetter, 1968). We, therefore, expect a larger number of irreparable breaks in glu and gly, and it is likely that these breaks lead to high SOS induction before cell death. In other words, the cells that die with a high-SOS phenotype likely die because of an irreparable DSB. Treatment with TET, at the low concentration that we used, would not change these cells’ replication status, therefore explaining the lack of suppression of the CIP-TET combination in this sub-population.

The second population of cells that died under CIP exposure did not exhibit a level of SOS higher than that of the cells that survived. Nevertheless, they stopped cell elongation and protein production. Contrary to the high-SOS population, these cells were more abundant in fast growth (glu-aa) than in glu and gly and their frequency diminished markedly when exposed to the CIP-TET antibiotic combination. This indicates that the suppressive effect of CIP-TET is mostly driven by the increased survival of these low SOS cells. Because of their limited SOS induction and higher abundance in glu-aa (when a large part of the chromosome is present in multiple copies because of multi-fork replication), we hypothesise that cellular death is not due to an irreparable break. This suggests that these cells die through a another molecular process that has yet to be fully characterized. To better understand their characteristics, we computed the elongation rate over a two-hour period before death of the low-SOS cells, the high-SOS cells and the cells that survived (for which we used a 2-hour period during antibiotic exposure). In all growth conditions, the elongation rate of the low-SOS cells that died was higher on average than that of the cells that survived or induced high SOS levels (Supplementary Figure S10, left panel). However, this difference disappeared in the CIP-TET condition suggesting that the reduction in elongation rate due to TET may underlie the better survival of these cells. This effect was stronger in glu-aa than in glu and gly. This is expected because we observed a growth-dependent effect of TET on cell elongation: the elongation rate of individual cells treated with TET decreased to a greater extent in glu-aa than in glu and gly (Supplementary Table S1) similar to what has been documented previously at the population level (Scott et al., 2010; Greulich et al., 2015). These results suggest that the death of the low-SOS cells is linked to a too-high elongation rate, which is compensated for by the reduction of elongation due to TET treatment. It is possible that cells that elongate too fast are not able to down-regulate ribosomal expression upon ciprofloxacin treatment, leading to cell death because of excess protein production as has been proposed previously based on population data (Bollenbach et al., 2009).

Whilst our observations help understand the growth dependence of the suppressive effect of TET on the low SOS cells, they do not fully elucidate the molecular mechanisms that lead to the death of these cells. Recent work suggests that cells that elongate transiently more slowly and induce increased levels of stress responses, such as acid resistance, survive better exposure to high levels of ciprofloxacin (Sampaio et al., 2022). However, we cannot directly link this phenomenon to our observations: in our case, elongation reduction by TET treatment is expected to also lead to a reduction of protein production and, therefore, limit the induction of various stress responses. Indeed, we observe that the increase in survival in CIP-TET for the low-SOS cells is not due to an increase in SOS expression: as expected from proteome global constraints, the SOS response is reduced in gly and glu under CIP-TET compared to CIP treatment and does not change in glu-aa (Figure 4A-C). Alternatively, it has been proposed that cell death under CIP exposure could be linked to formation of ROS compounds (Kohanski et al., 2007; Hong et al., 2020) or to an imbalance in energy metabolism where ATP becomes limiting for DNA repair (Chevereau and Bollenbach, 2015). Treatment with TET, which limits the high energy-consuming protein synthesis machinery, could free up ATP for DNA repair. It has also been shown that bacteriostatic antibiotics such as TET reduce cellular respiration, which results in blocking killing by bactericidal antibiotics such as CIP (Lobritz et al., 2015). Our results suggest that full elucidation of these mechanisms requires detailed quantification at the single-cell level using, for example, single-cell reporters of ATP concentration as has been done recently (Yaginuma et al., 2014; Lin and Jacobs-Wagner, 2022). In the longer term, a quantitative understanding of how individual cells respond to treatment can inform emergent population-level effects and improve the modelling of bacterial physiological responses to drug combinations.

## 5 Data availability

The data are currently available upon request and will be released on GitLab, Zenodo and BioImage archive upon manuscript publication.

## 7 Acknowledgements

James Broughton was supported by a PhD studentship from the Darwin Trust of Edinburgh. This work was supported by a Wellcome Trust Investigator Award (Grant No. 205008/Z/16/Z) to M.E.K. and by the United Kingdom Research and Innovation (grant EP/S02431X/1), UKRI Centre for Doctoral Training in Biomedical AI at the University of Edinburgh, School of Informatics (PhD studentship to Achille Fraisse). For the purpose of open access, the authors have applied a Creative Commons attribution (CC BY) licence to any author-accepted manuscript version arising.

## 8 Supplementary Figures and Tables

**Figure S1:**
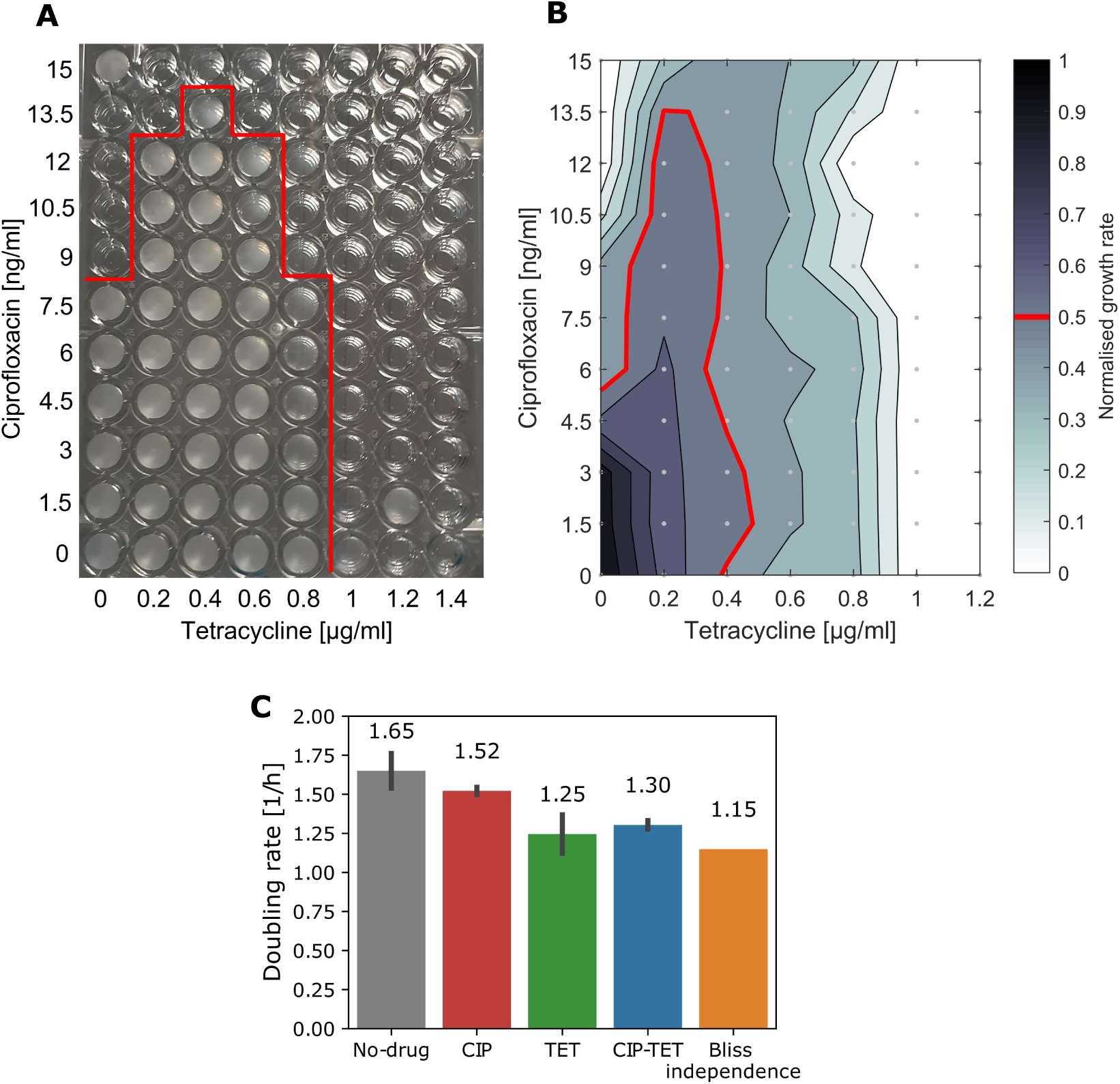
Bulk-level characterisation of the ciprofloxacin-tetracycline drug combination. **A-B**. Checkerboard assay showing suppressive interaction between ciprofloxacin and tetracycline. **A**. Picture of the 96-well plate indicating growth (turbidity) in wells after 28 hours incubation. Red line indicates wells with significant growth. Wells with significant growth outside this line are likely resistant mutants and were excluded from analysis. Cells were cultured in the glu-aa medium. **B**. Dose-response surface showing growth rates (blue colour) over a two-dimensional grid of antibiotic concentrations (grey dots represent concentrations used). Colour gradient represents normalised growth rate where dark blue represents no growth inhibition and white represents no growth. Red line follows the IC_50_ isobole on the dose-response surface. The bulging convex isobole is indicative of a suppressive drug interaction according to the Loewe additivity model (Loewe, 1928, 1953). Strain SJR206 was used for the checkerboard assay. **C**. Bulk doubling rates under 3 ng/ml ciprofloxacin and 0.2 *µ*g/ml tetracycline mono-exposure and under combination treatment in the glu-aa growth medium. Shown is the mean (value indicated above each column) and standard deviation from at least three experimental replicates. The additive prediction for the doubling rate under the antibiotic combination was calculated according to the Bliss definition of independent drug interactions (Bliss, 1939) which assumes that individual drug effects can be multiplied: *g_AB_* = *g_A_ × g_B_*, where *g_X_* is the normalised doubling rate in the presence of antibiotic X. Finally, *g_AB_* was multiplied by the doubling rate of the no-drug control to get the Bliss additive prediction. Since the doubling rate under CIP-TET is higher than the additive expectation, this indicates an antagonistic drug interaction. Doubling rates were measured using automatic OD_600_ measurements from OGI-BIO bioreactors for strain JKB43. Detailed description of protocols can be found in the Supplementary Methods.

**Figure S2:**
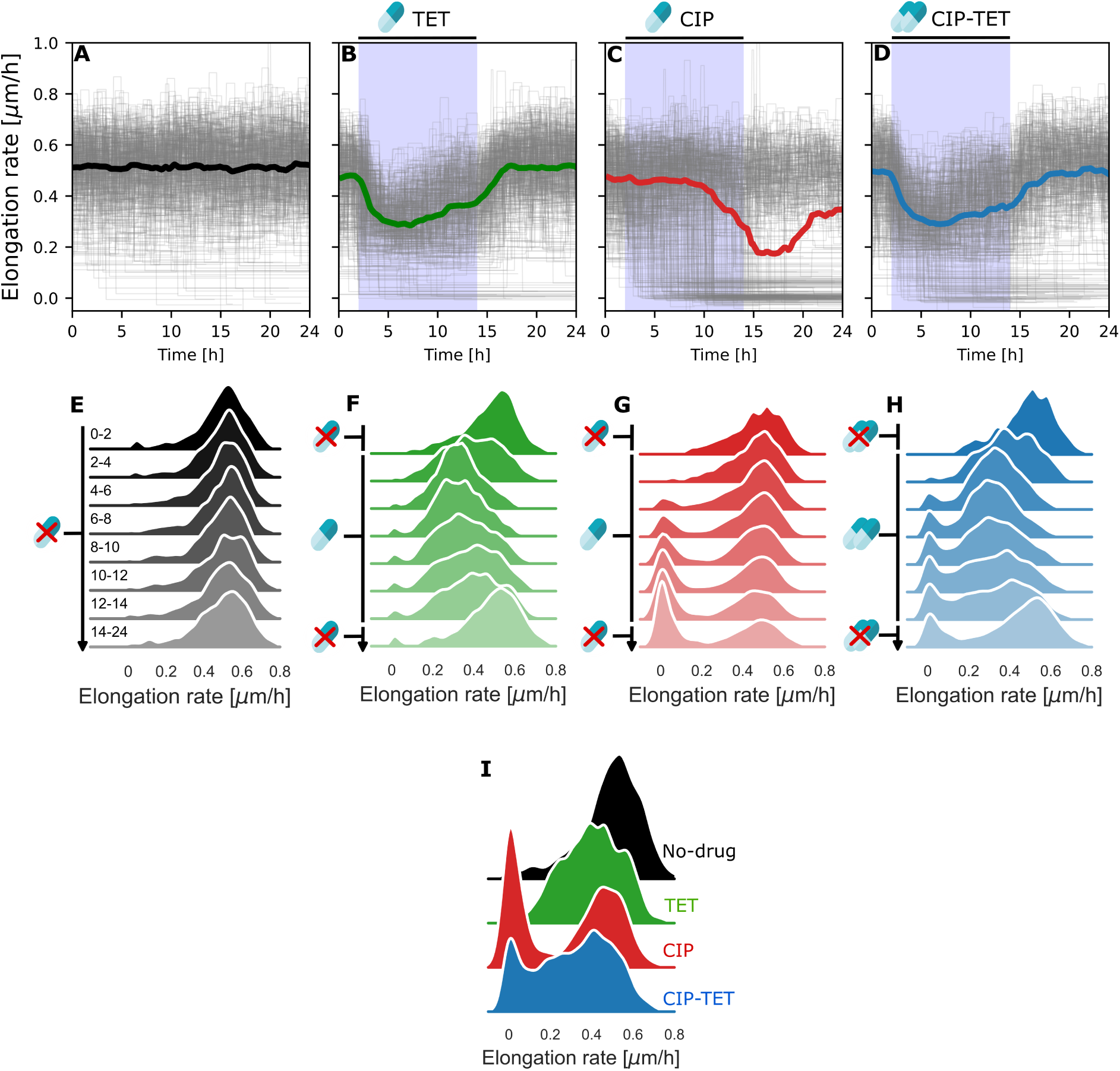
Single-cell elongation rates under sub-lethal antibiotic treatment for growth in the glu medium. **A-D**. Single-cell trajectories of elongation rates under no-drug (A, n = 342 cells), 0.2 *µ*g/mL tetracycline (B, n = 351), 3 ng/mL ciprofloxacin (C, n = 386), and CIP-TET (D, n = 355) treatment. Antibiotics were introduced between hours 2-14, indicated by the blue shaded area. Grey lines represent individual mother cell lineage trajectories. Solid lines represent the median of the population. Shown is data from one experimental replicate for growth in the glu medium. **E-H**. Distributions of single-cell elongation rates for the no-drug control (E), TET treatment (F), CIP treatment (G), and CIP-TET treatment (H) drawn from sequential two-hour periods under different antibiotic treatments. Distributions are kernel density estimates of the underlying histogram pooled from at least two experimental replicates. Time periods, in hours, are indicated to the left of each distribution. Antibiotics were introduced from hours 2-14 as indicated. **I**. Distributions of single-cell elongation rates drawn from the final 2 hours of antibiotic treatment.

**Figure S3:**
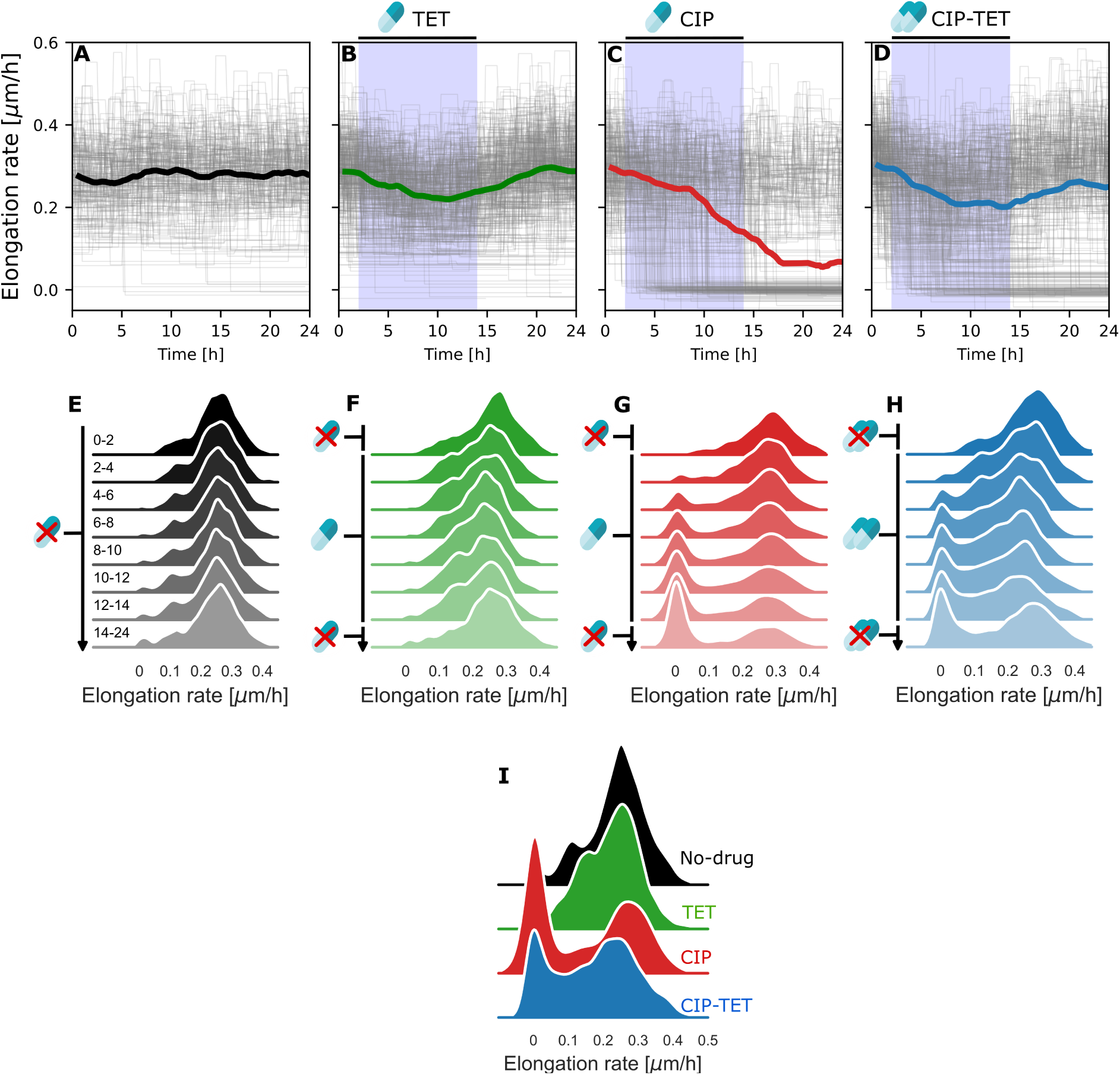
Single-cell elongation rates under sub-lethal antibiotic treatment for growth in the gly medium. **A-D**. Single-cell trajectories of elongation rates under no-drug (A, n = 188 cells), 0.2 *µ*g/mL tetracycline (B, n = 317), 3 ng/mL ciprofloxacin (C, n = 252), and CIP-TET (D, n = 328) treatment. Antibiotics were introduced between hours 2-14, indicated by the blue shaded area. Grey lines represent individual mother cell lineage trajectories. Solid lines represent the median of the population. Shown is data from one experimental replicate for growth in the gly medium. **E-H**. Distributions of single-cell elongation rates for the no-drug control (E), TET treatment (F), CIP treatment (G), and CIP-TET treatment (H) drawn from sequential two-hour periods under different antibiotic treatments. Distributions are kernel density estimates of the underlying histogram pooled from at least two experimental replicates. Time periods, in hours, are indicated to the left of each distribution. Antibiotics were introduced from hours 2-14 as indicated. **I**. Distributions of single-cell elongation rates drawn from the final 2 hours of antibiotic treatment.

**Figure S4:**
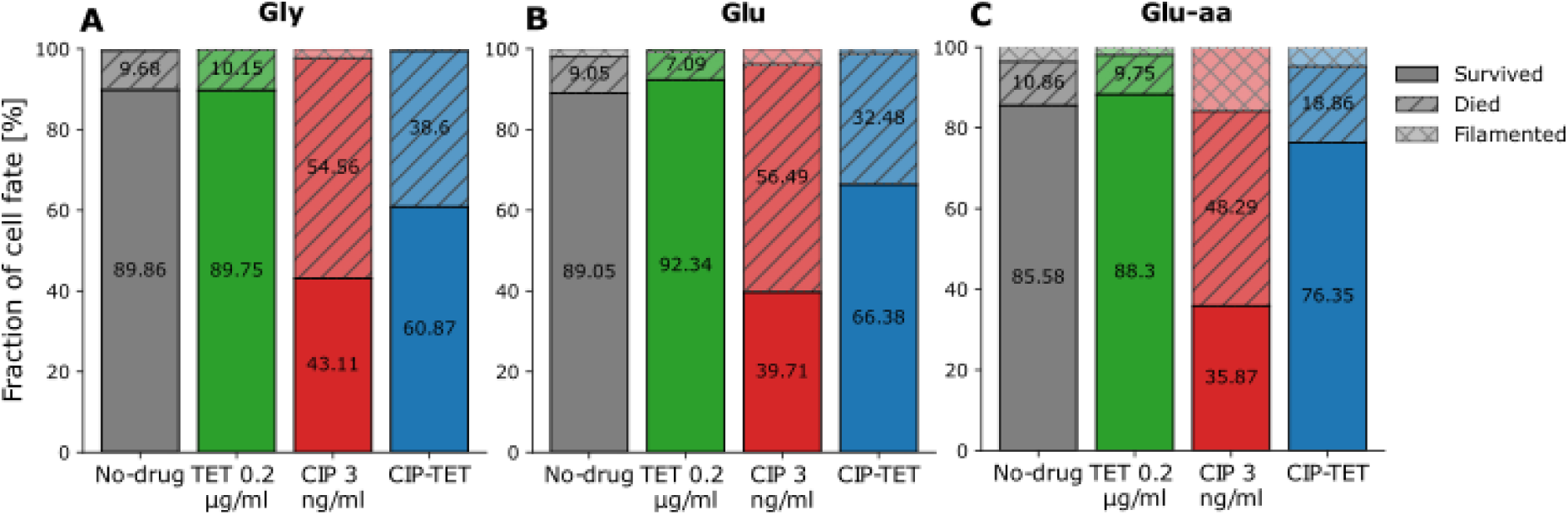
Suppression of cell death under the CIP-TET combination is growth-dependent. **A**. Proportion of cells that either survived, died (growth and division arrest or lysis), or filamented under different treatment and growth conditions over the duration of the experiment.

**Figure S5:**
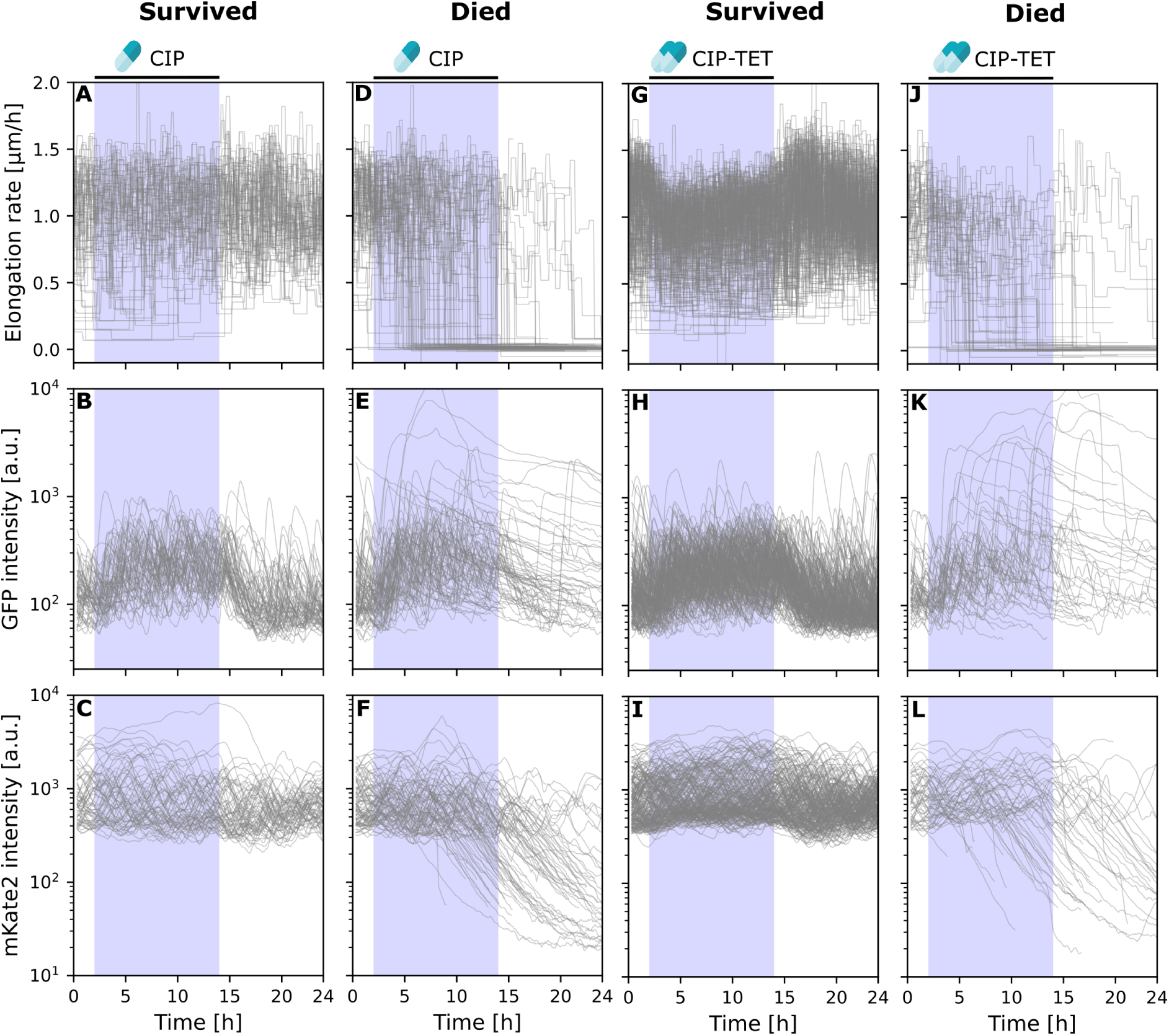
Single-cell responses separated by cell fate for the glu-aa growth medium. Lineage fate at the end of the experiment was classified as ‘survived’ or ‘died’ as described previously. **A-F**. Single-cell elongation rates, SOS expression, and constitutive expression under CIP treatment. **G-L**. Single-cell elongation rates, SOS expression, and constitutive expression under CIP-TET treatment.

**Figure S6:**
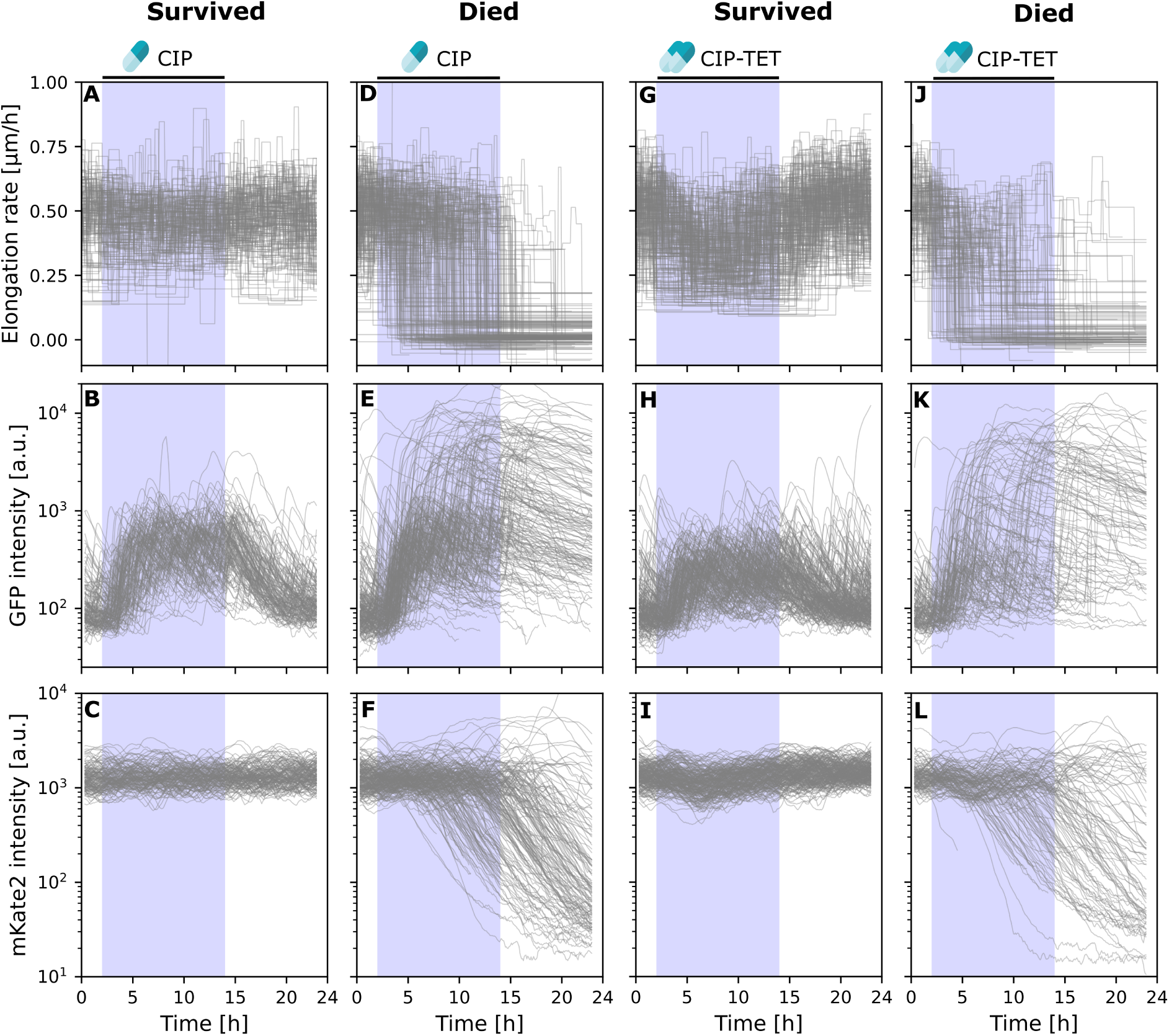
Single-cell responses separated by cell fate for the glu growth medium. Lineage fate at the end of the experiment was classified as ‘survived’ or ‘died’ as described previously. **A-F**. Single-cell elongation rates, SOS expression, and constitutive expression under CIP treatment. **G-L**. Single-cell elongation rates, SOS expression, and constitutive expression under CIP-TET treatment.

**Figure S7:**
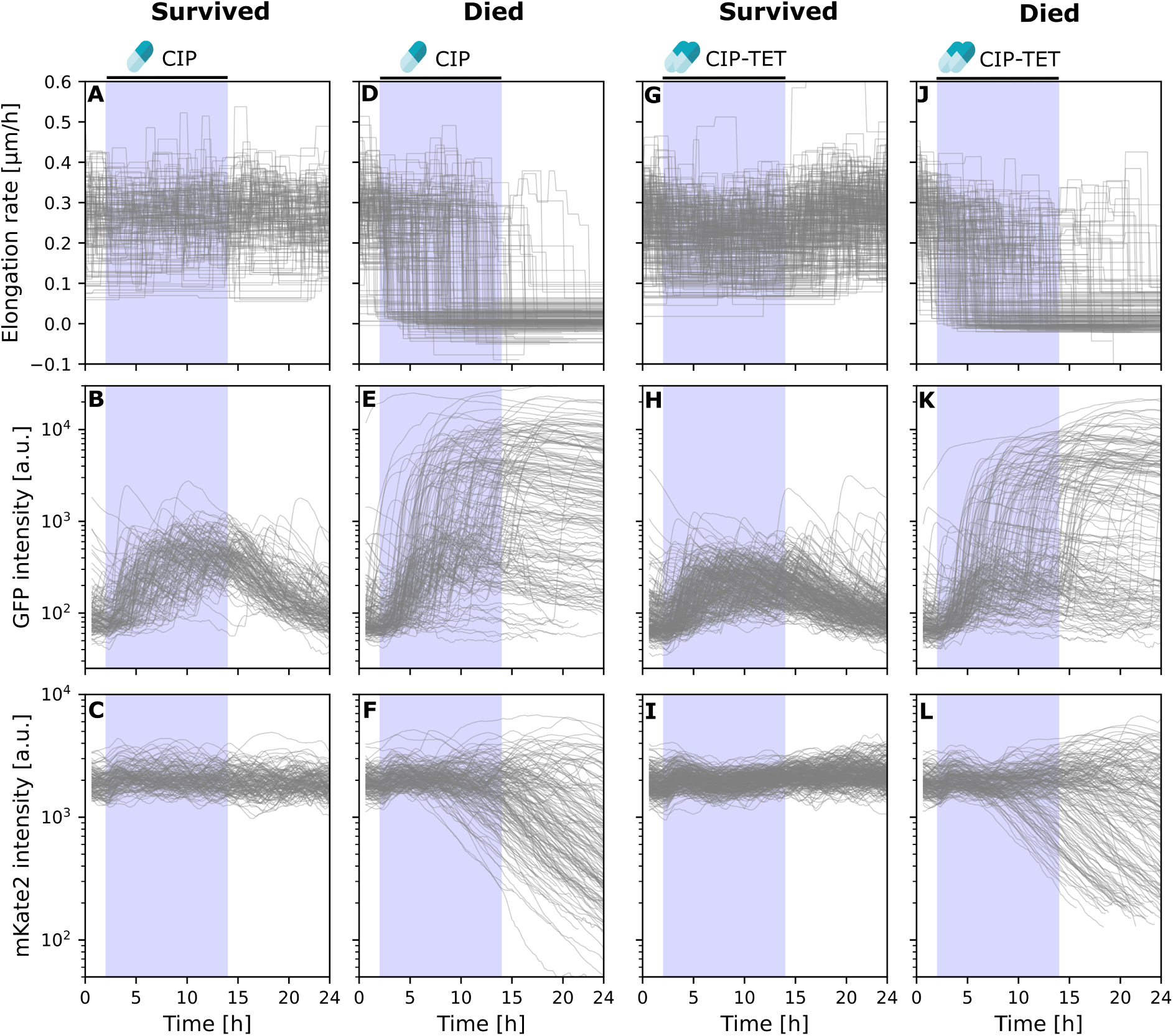
Single-cell responses separated by cell fate for the gly growth medium. Lineage fate at the end of the experiment was classified as ‘survived’ or ‘died’ as described previously. **A-F**. Single-cell elongation rates, SOS expression, and constitutive expression under CIP treatment. **G-L**. Single-cell elongation rates, SOS expression, and constitutive expression under CIP-TET treatment.

**Figure S8:**
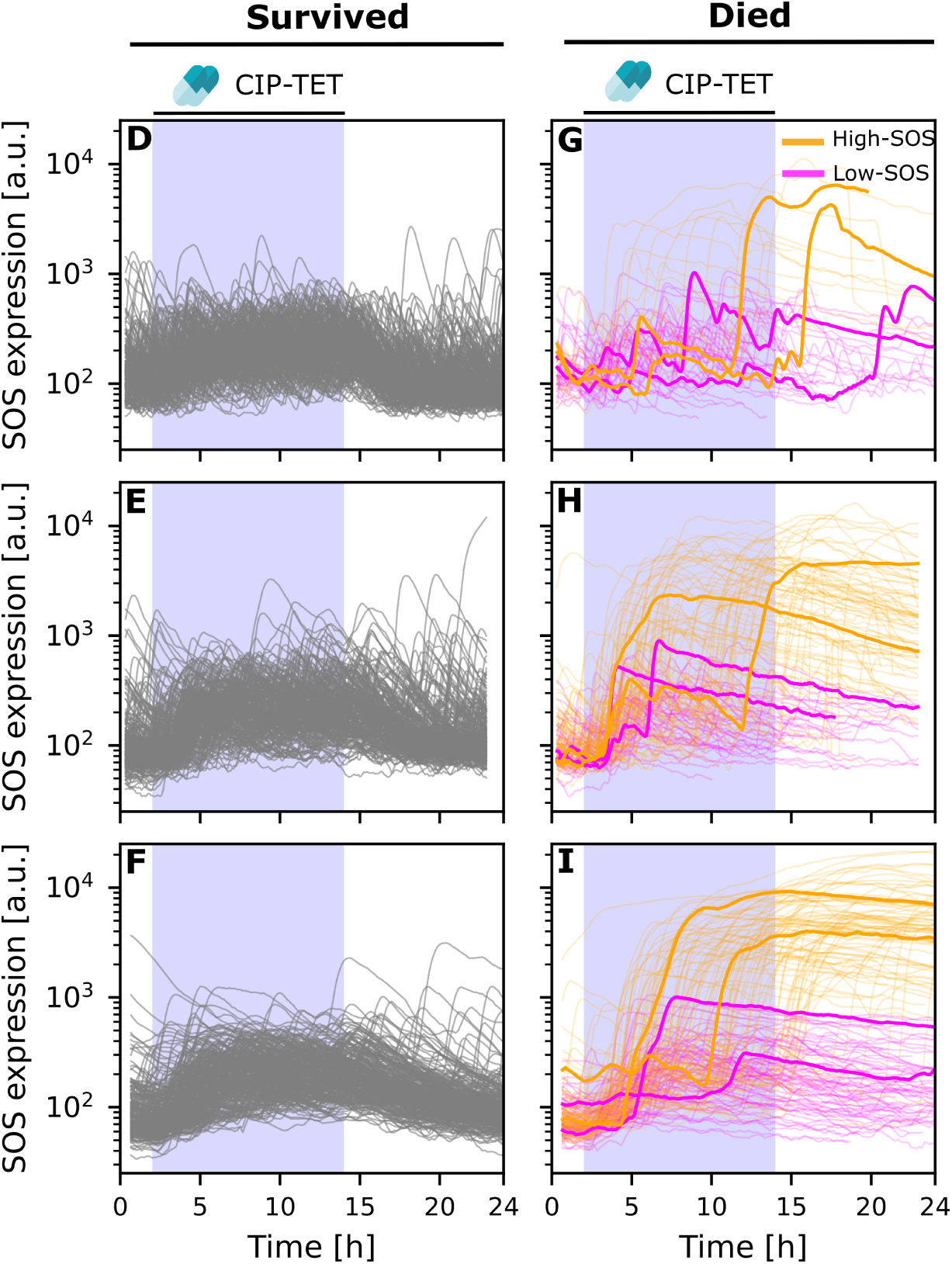
The SOS response is highly heterogeneous under CIP-TET treatment. **A**. Single cell trajectories of SOS expression classified by expression level (low-SOS: magenta and high-SOS: orange) for dead cells under CIP-TET treatment in different growth conditions. Example trajectories are highlighted to illustrate the behaviour of the two sub-populations. Shown is data from one experimental replicate for clarity.

**Figure S9:**
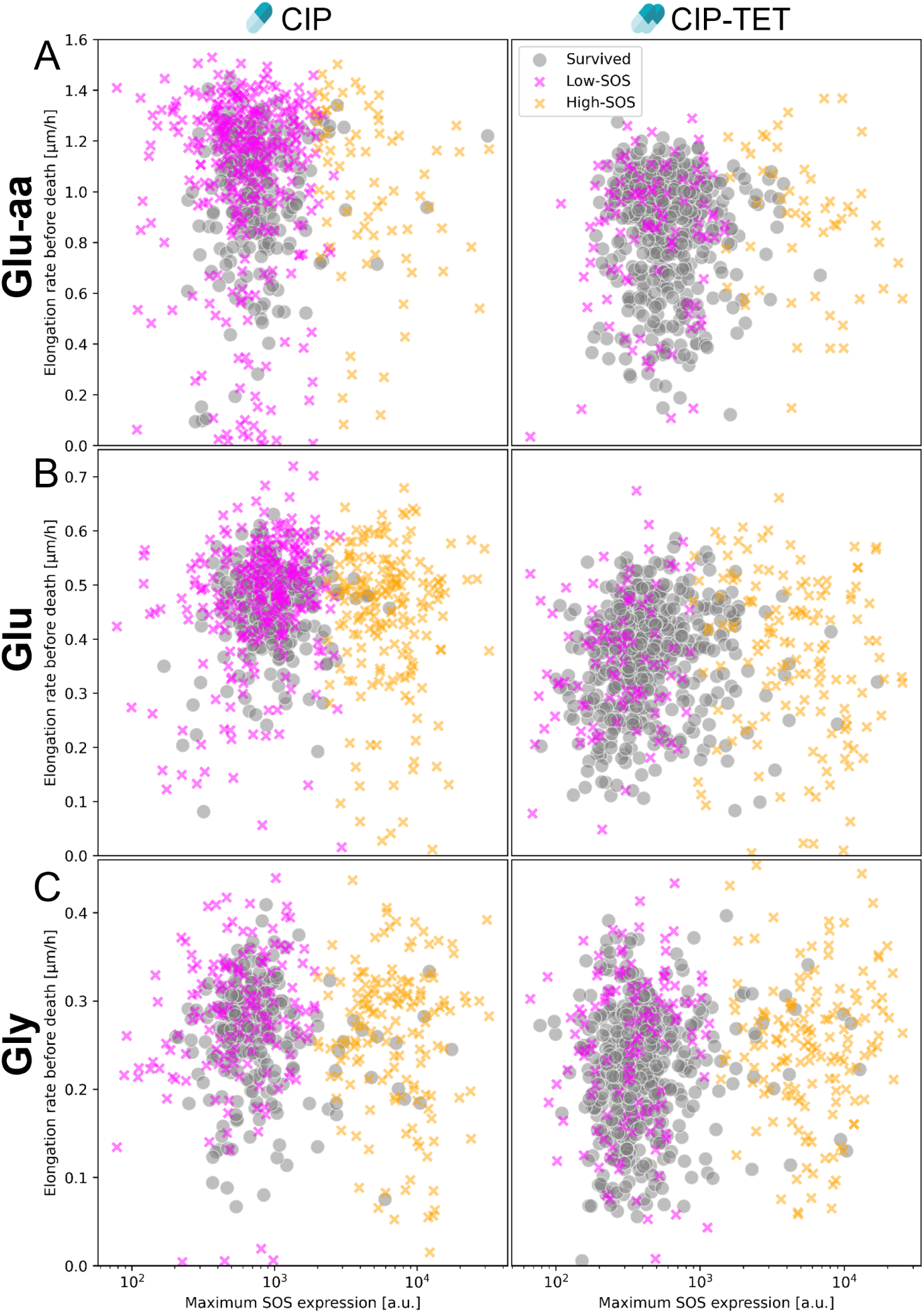
Low-SOS cells are protected by treatment with under the combination of antibiotics to a greater extent than high-SOS cells. The average elongation rate in the two hours preceding death for dead cells and in the second half of the antibiotic exposure period for survivors is plotted against the maximum SOS intensity over the whole trajectory.

**Figure S10:**
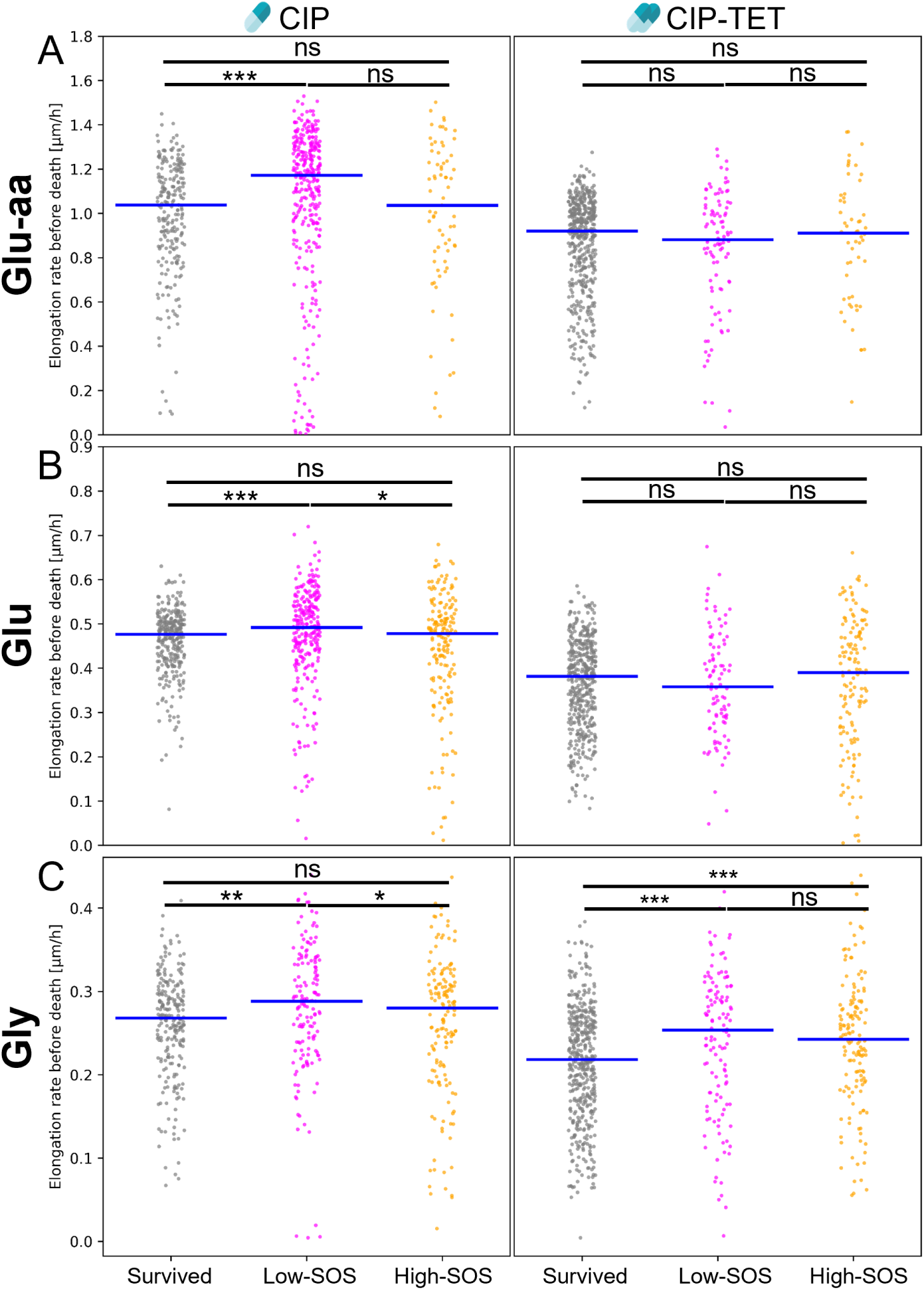
Low-SOS dying cells grow faster during the two hours before they die in the CIP experiment. Data points represent the average of single-cell elongation rates in the two hours preceding death for dead cells (low-SOS and high-SOS). For surviving cells, we show the average of singlecell elongation rates over the second half of the antibiotic exposure period which represents a steady-state response. The median of the population is indicated by the blue bar. Results of Mann-Whitney tests are shown above (ns: non significant, **\***: *p <*0.05, **\*\***: *p <*0.01, **\*\*\***: *p <*0.001.

**Table S1:** Single-cell statistics under different treatment conditions for experimental replicates in different growth media. Data shown is representative of data pooled over the duration of the experiment for the no-drug control, or over the final 2 hours of antibiotic treatment.

| Treatment | Medium<br>& replicate | Number<br>lineages | Number<br>cell-cycles | Elongation rate [$\mu\text{m/h}$] | | | Division time [min] | | | SOS intensity<br>[a.u./area] | | | Constitutive intensity<br>[a.u./area] | | |
| --- | --- | --- | --- | --- | --- | --- | --- | --- | --- | --- | --- | --- | --- | --- | --- |
|  |  |  |  | Mean | Median | CV | Mean | Median | CV | Mean | Median | CV | Mean | Median | CV |
| No-drug<br>control | Glu<sub>aa</sub>_1 | 283 | 8919 | 1.09 | 1.13 | 0.27 | 40.54 | 35 | 0.47 | 152.61 | 115.61 | 1.18 | 791.09 | 626.33 | 0.6 |
|  | Glu<sub>aa</sub>_2 | 334 | 11581 | 1.09 | 1.12 | 0.27 | 39.74 | 35 | 0.53 | 140.51 | 107.72 | 1.2 | 683.03 | 531.85 | 0.63 |
|  | Glu_1 | 78 | 1108 | 0.51 | 0.52 | 0.25 | 81.12 | 80 | 0.32 | 151.25 | 98.35 | 1.48 | 1350.84 | 1312.4 | 0.21 |
|  | Glu_2 | 342 | 5871 | 0.53 | 0.54 | 0.23 | 83.97 | 75 | 0.39 | 150.19 | 95.59 | 2.55 | 1362.13 | 1306.9 | 0.25 |
|  | Gly_1 | 188 | 1978 | 0.28 | 0.28 | 0.29 | 166.23 | 150 | 0.47 | 119.2 | 79.39 | 2.56 | 1942.07 | 1928.56 | 0.24 |
|  | Gly_2 | 376 | 3517 | 0.25 | 0.26 | 0.28 | 191.02 | 170 | 0.5 | 149.37 | 88.68 | 2.46 | 2070.4 | 2028.95 | 0.17 |
|  | Gly_3 | 304 | 2647 | 0.24 | 0.25 | 0.29 | 204.31 | 180 | 0.57 | 167.08 | 96.46 | 2.8 | 2130.1 | 2093.16 | 0.17 |
| CIP | Glu<sub>aa</sub>_1 | 189 | 511 | 0.89 | 1.04 | 0.53 | 45.73 | 40 | 0.77 | 337.56 | 235.44 | 2.24 | 621.8 | 508.2 | 0.81 |
|  | Glu<sub>aa</sub>_2 | 156 | 423 | 0.89 | 1.02 | 0.51 | 46.27 | 40 | 0.57 | 289.77 | 226.37 | 1.01 | 638.85 | 495.08 | 0.79 |
|  | Glu<sub>aa</sub>_3 | 182 | 522 | 0.95 | 1.11 | 0.52 | 42.57 | 35 | 0.76 | 338.52 | 219.22 | 1.76 | 576.58 | 420.37 | 0.83 |
|  | Glu_1 | 379 | 678 | 0.36 | 0.43 | 0.61 | 101.39 | 85 | 0.86 | 893.88 | 389.61 | 2.33 | 1140.08 | 1109.73 | 0.51 |
|  | Glu_2 | 357 | 620 | 0.34 | 0.41 | 0.65 | 109.6 | 90 | 0.8 | 1133.67 | 519.67 | 1.75 | 1171.14 | 1182.86 | 0.46 |
|  | Gly_1 | 251 | 362 | 0.18 | 0.21 | 0.78 | 222.51 | 160 | 0.98 | 1402.11 | 386.83 | 2.28 | 1958.16 | 1882.28 | 0.33 |
|  | Gly_2 | 307 | 442 | 0.17 | 0.21 | 0.82 | 199.46 | 150 | 0.81 | 1598.73 | 451.36 | 1.91 | 1948.16 | 1898.42 | 0.34 |
| TET | Glu<sub>aa</sub>_1 | 320 | 1079 | 0.91 | 0.96 | 0.32 | 54.43 | 45 | 0.73 | 135.06 | 106.78 | 0.96 | 1130.22 | 862.97 | 0.64 |
|  | Glu<sub>aa</sub>_2 | 233 | 793 | 0.92 | 0.96 | 0.3 | 50.77 | 40 | 0.65 | 134.29 | 100.25 | 1.25 | 992.38 | 765 | 0.66 |
|  | Glu_1 | 349 | 714 | 0.38 | 0.4 | 0.34 | 118.86 | 100 | 0.57 | 162.74 | 88.29 | 2.59 | 1381.54 | 1329.35 | 0.22 |
|  | Glu_2 | 351 | 791 | 0.46 | 0.48 | 0.3 | 101.22 | 85 | 0.5 | 157.16 | 95.57 | 3.2 | 1416.62 | 1362.65 | 0.24 |
|  | Gly_1 | 317 | 534 | 0.23 | 0.23 | 0.39 | 202.62 | 170 | 0.54 | 123.91 | 77.37 | 2.82 | 2154.14 | 2142.47 | 0.19 |
|  | Gly_2 | 277 | 449 | 0.2 | 0.21 | 0.35 | 229.67 | 190 | 0.59 | 146.71 | 89.56 | 2.25 | 2212.07 | 2204.12 | 0.16 |
|  | Gly_3 | 411 | 684 | 0.23 | 0.25 | 0.35 | 205.81 | 160 | 0.69 | 280.5 | 97.44 | 4.31 | 2195.34 | 2101.36 | 0.23 |
| CIP-TET | Glu<sub>aa</sub>_1 | 320 | 1004 | 0.87 | 0.93 | 0.36 | 53.33 | 45 | 0.59 | 243.99 | 191.94 | 1.61 | 935.18 | 720.29 | 0.69 |
|  | Glu<sub>aa</sub>_2 | 287 | 935 | 0.92 | 0.98 | 0.34 | 49.75 | 45 | 0.53 | 301.3 | 214.66 | 1.68 | 891.93 | 684.24 | 0.66 |
|  | Glu_1 | 347 | 653 | 0.37 | 0.4 | 0.46 | 121.59 | 100 | 0.8 | 430.8 | 194.92 | 2.45 | 1326.25 | 1305 | 0.36 |
|  | Glu_2 | 336 | 598 | 0.34 | 0.36 | 0.56 | 134.51 | 105 | 0.78 | 588.37 | 229 | 2.04 | 1280.3 | 1254.59 | 0.38 |
|  | Gly_1 | 325 | 503 | 0.2 | 0.23 | 0.65 | 212.31 | 160 | 0.69 | 754.01 | 205.6 | 2.4 | 2194.44 | 2135.45 | 0.32 |
|  | Gly_2 | 431 | 634 | 0.17 | 0.19 | 0.59 | 237.86 | 190 | 0.67 | 703.59 | 219.16 | 2.19 | 2027.37 | 2043.05 | 0.26 |

## 9 Supplementary Methods

### 9.1 List of strains, plasmids, and primers

Here are the list of strains, plasmids, and primers used in this study. Details for the construction of the parental strain eSJR206 can be found in Jaramillo-Riveri et al. (2022). Bacterial strains were constructed by P1 transduction and the resistance markers removed using the pE-FLP plasmid (St-Pierre et al., 2013). After construction, all strains were checked by PCR amplification and Sanger sequencing of the modified chromosomal region. The *motA* gene was deleted to reduce motility and improve retention of cells in the mother machine microchannels.

**Table S2:** List of strains. P1 stands for P1 phage transduction.

| Strain | Background | Genotype | Source/Construction |
| --- | --- | --- | --- |
| JW1879-2 | BW25113 | *rph-1* $\lambda^-$ *F*-<br>*hsdR514*<br>$\Delta(araD-araB)567$<br>$\Delta(rhaD-rhaB)568$<br>$\Delta lacZ4787(::rrnB-3)$<br>$\Delta motA743::Kn^R$ | Gift from Rosalind Allen (CGSC 9565) (Baba et al., 2006) |
| eSJR206 | MG1655 | *rph-1* $\lambda^-$ *F*-<br>*HK022*:P<sub>*sulA*</sub>-*mGFP*<br>*P21*:P<sub>*tet01*</sub>-*mKate2* | See Jaramillo-Riveri et al. (2022) for construction |
| JKB22 | MG1655 | eSJR206<br>$\Delta motA::Kn^R$ | eSJR206 P1 using JW1879-2 |
| JKB43 | MG1655 | eSJR206<br>$\Delta motA$ | JKB22 pE-FLP using pSJR017 |

**Table S3:**
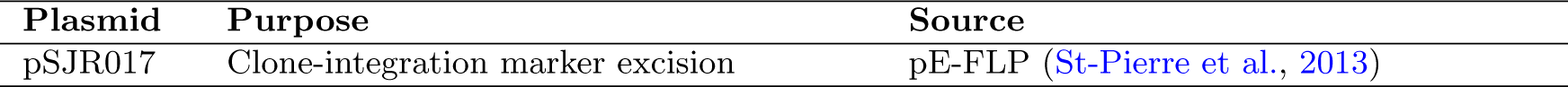
List of plasmids.

**Table S4:**
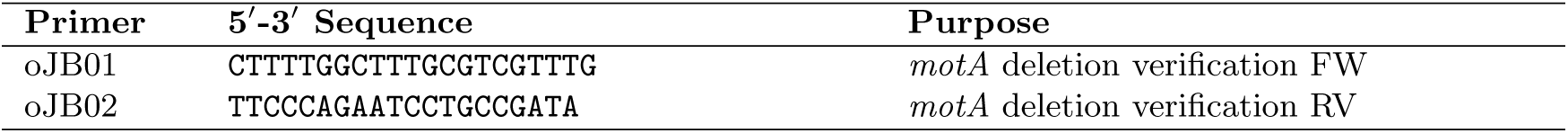
List of primers.

### 9.2 Bulk experiments

#### 9.2.1 Checkerboard assay

The inoculum was prepared as follows: An LB overnight culture was prepared as described in the Materials and Methods. Cells were then diluted (1:2000) into 5 mL of M9-based medium and incubated overnight at 37°C with agitation (150 rpm). In the morning, another 1:200 dilution was performed and the culture was incubated until OD_600_ = 0.2. In flat-bottomed 96-well plates (Corning Costar), a concentration series for each antibiotic was set up in all columns (drug A) or rows (drug B) as follows: Antibiotics were made up at double the highest concentration desired in 5 mL M9-based medium. A gradient dilution was set up on the x-axis by adding 100 *µ*L of drug A to all wells in the first column and a serial dilution was performed for the remaining columns of wells up to column 10, with no drug added to column 11 (Supplementary Figure S11). Thus, each well was made up to a volume of 100 *µ*L with double the desired drug concentration. A second drug gradient was similarly set up across the y-axis (rows) for drug B, again at double the desired concentration, with no drug added to row H. The desired concentration was achieved in drug combination wells when the two antibiotic solutions are added (100 *µ*L drug A + 100 *µ*L drug B = 1:2 dilution). For wells administered with a single drug (column 11 and row H), 100 *µ*L sterile medium was added to make up the final concentration (1:2 dilution). All wells were then mixed using a pipette. No drugs were added to column 12 which contained the positive and negative controls. At one edge of the plate (column H), four wells were filled with 200 *µ*L growth medium and inoculum (positive growth controls) and the other four with just 200 *µ*L growth medium (negative controls for contamination). All wells of the plate (except the negative controls) were inoculated at a 1:1000 dilution using the prepared culture described above. All solutions were pre-warmed at 37°C.

**Figure S11:**
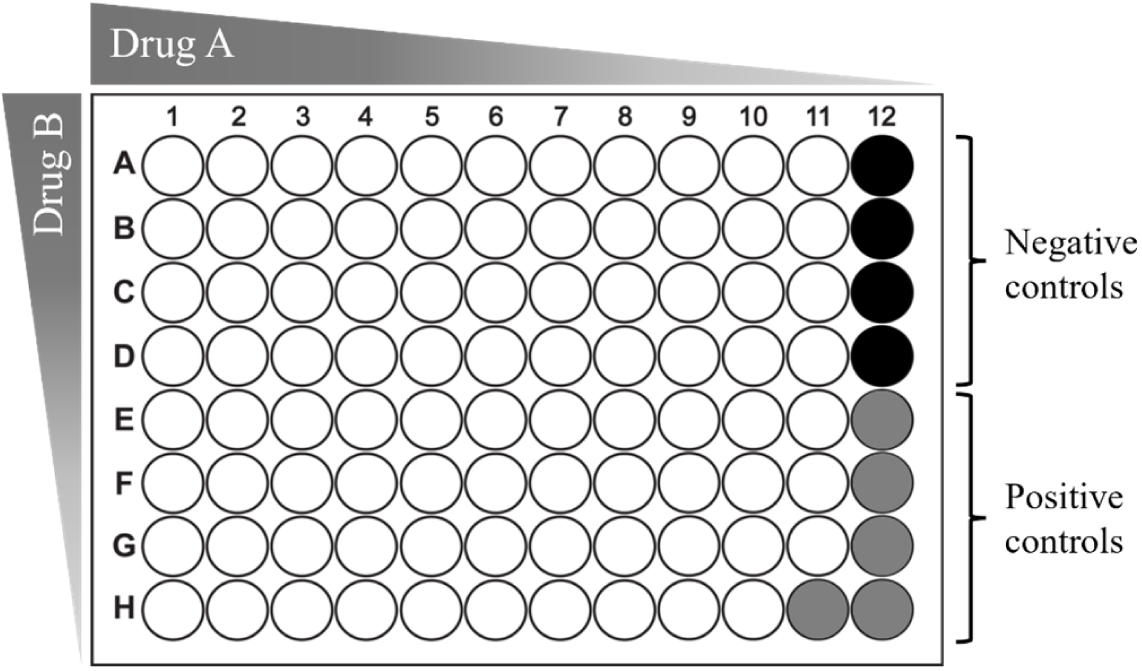
Depiction of checkerboard assay in a 96-well plate used to evaluate interactions for antibiotic combinations. A 2D concentration gradient is set up for drug A and drug B along the X and Y axis. Single drug concentration gradients are set up in Row H and Column 11 for drug A and B, respectively. The positions of the negative (black wells) and positive controls (grey wells) are indicated.

A lid was taped onto the 96-well plate to minimise evaporation and debris formation (the tape prevents movement of the lid during shaking). The 96-well plate was then inserted into the FLUOstar Omega microplate reader (BMG LABTECH) and OD_600_ was measured for each well every 7 min for 28-50 hours depending on the growth medium. The platereader maintained a temperature at 37°C and shaking at 700 rpm using the double-orbital shaking mode. All ODs were blank corrected (subtracted by initial OD of sterile medium) and then exported to an Excel spreadsheet and imported to MATLAB for further analysis.

Growth curves from the platereader were noisy and sometimes contained extreme values considered to be measurement errors. Therefore, growth curves were smoothed with a 5-window moving median and outliers corrected using the *filloutliers* function in MATLAB. Outliers that were missed were manually corrected. Growth rates in each well were then measured via linear regression (*regress* function) to log-transformed OD_600_ from the exponential phase of the growth curve (between 0.025 ¡ OD_600_ ¡ 0.25). Fits with an R^2^ value below 0.8 were discarded. Growth rates were then normalised by the control growth rate (averaged from five wells). To reduce noise, a cubic smoothing spline was used for interpolation to the measured growth rates using the *csaps* MATLAB function. The dose response surface (linearly interpolated isoboles) was then plotted using the *contour* function in MATLAB.

#### 9.2.2 Bulk growth rates

Population (bulk) growth rates were measured automatically using OGI-BIO bioreactors. The bioreactor was was calibrated according to the manufacturer instructions. The inoculum was prepared as described previously for the checkerboard assay. Tubes containing 10 ml of M9-based medium were then inoculated at a 1:1000 dilution using the prepared culture. The bioreactor was incubated at 37°C for 24 hours. The cultures were mixed at 2000 rpm by a magnetic stir bar. OD measurements were taken every 5 minutes. The excel output file containing the OD measurements were then analysed in MATLAB. Growth rates were estimated via linear regression to the log-transformed growth curves during the exponential growth phase. Doubling rates (*λ*) were converted from growth rates (*µ*) as: *λ* = *ln*(2) *× µ*.

### 9.3 Mother machine

#### 9.3.1 Microfluidic device dimensions

**Table S5:**
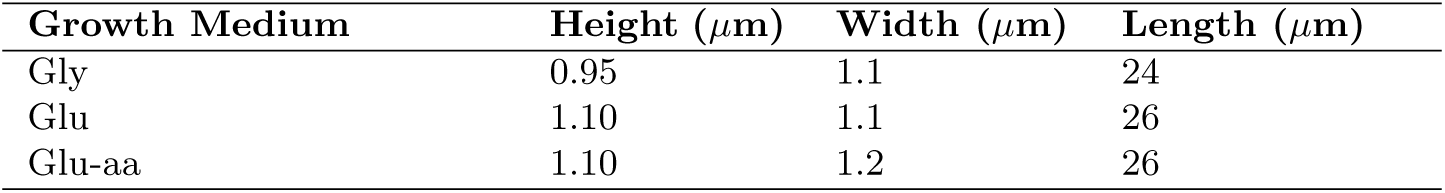
Mother machine microchannel dimensions used for different growth media.

| Growth Medium | Height ($\mu\text{m}$) | Width ($\mu\text{m}$) | Length ($\mu\text{m}$) |
| --- | --- | --- | --- |
| Gly | 0.95 | 1.1 | 24 |
| Glu | 1.10 | 1.1 | 26 |
| Glu-aa | 1.10 | 1.2 | 26 |

#### 9.3.2 Classification of cell fate

Using a simple algorithm, we classified cells into three categories by whether they continued growth and division (‘survived’), abruptly stopped growth and division (‘died’), or excessively filamented beyond the length of the microchannels. We quantified the fate of mother cell lineages over the duration of the experiment (24 hours), which included a 10 hour recovery period. This ensures that cells initially classified as ‘dead’ were not persister cells (i.e. resumed growth post antibiotic treatment). First, lineages that excessively filamented (length *<*20 *µ*m) were separated and classified as ‘hyper-filamented’. These cells extended beyond the length of the microfluidic microchannels (cell traps). The remaining lineages were classified as ‘died’ if their last recorded elongation rate is below a threshold value, or if they did not divide for 4 hours or more. The elongation rate threshold was set at 20% of the median elongation rate of the no-drug control condition for each growth medium. In some rare cases cells lysed – immediately losing their physical integrity – and could no longer be visually tracked, therefore escaping the filters set above. These cells were visually identified and manually classified as ‘died’.

The majority of cells that ceased growth maintained physical integrity and did not resume growth after removal of antibiotic treatment. These cells exhibited, after a delay, an exponential decline in mKate2 intensity (constitutive gene expression) once growth had stopped and did not recover (Supplementary Figure S5-S7F & L). Given that elongation (dilution) had ceased, this decline indicates that constitutive expression and metabolic activity/protein production has stopped completely. The exponential decline in mKate2 intensity is likely due to photobleaching of the fluorescent protein. This observation further suggests that these cells are most likley dead or in the process of dying. We can therefore use the constitutive expression trajectories to verify the classification of dead cells. An example of mother cell lineages classified by fate is shown in Supplementary Figures S5-S7. There was good separation between ‘survived’ and ‘dead’ lineages based on the elongation rates and constitutive gene expression trajectories.

#### 9.3.3 Quantification of cell survival and death

Survival analysis was carried out using the the Kaplan-Meier (K-M) estimator (Kaplan and Meier, 1958), a non-parametric model commonly used for survival analysis. First, we organised our data into two columns, with the first column populated with lineage lifespans (survival times) in hours, and the second column denoting whether a death event occurred or not (Boolean, true or false). Not all cells died during the experiment – these are known as right-censored individuals and are important to include in our statistic to avoid overestimating the rate of death. Lifespans were determined from the start of the experiment until the time of death or time of censorship, where censorship refers to the end of the experiment or until lineage tracking is lost. For lineages classified as ‘dead’ (see above), the time of death was determined as the start of the cell-cycle during which cell death occurred (i.e. time of birth of the dead cell). Each lineage is made up of a succession of cell-cycles. Here, a cell-cycle represents the period between birth and division or between birth and the last recorded image. A dead cell therefore represents a lineage’s final cell-cycle. Although we could not precisely determine the fate of lineages that escaped due to filamentation, we lumped them with dead lineages in the survival curves since they are not growing ‘normally’ and to avoid bias if these were removed from the calculation. The survival function S(t) was estimated using the K-M model (Kaplan and Meier, 1958), defined as:

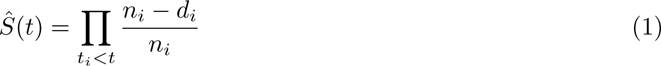

where *d_i_* are the number of death events at time *t* and *n_i_* is the number of subjects at risk of death just prior to time *t*. The K-M estimate represents an accumulation of probabilities equivalent to *S(t)* = P(surviving past time *t*). The analysis was implemented using the lifelines package in Python (Davidson-Pilon, 2019).

#### 9.3.4 Classification of dead lineages into low-SOS and high-SOS inducers

The cells that died during the experiment (fate classification described above) were split into two groups based on their SOS expression. The “high inducer” cells undergo a jump in SOS expression correlated to the moment they stop elongating (for example see Supplementary Figure S6D & E) while “low inducers” stop expressing SOS and their fluorescence simply decay (both group can be seen in S8G-I). To detect such groups we applied a threshold on the maximum SOS value of each trajectory, since high inducers present a fluorescence peak. Trajectories whose maximum value are above the threshold were classified as high inducer while other trajectories were classified as low inducers (see the two populations in yellow and purple respectively in Supplementary Figure S9. This value was set independently for each condition and replicate. A density plot of the maximum values of each trajectory often show two distinct distributions for the high and low SOS inducers (see Supplementary Figure S9) and the threshold value was set to the valley between those populations when possible, with additional manual adjustments when needed. The list of thresholds for each condition and replicate are shown in Supplementary Table S6.

**Table S6:** Thresholds log values used for classification of low and high SOS inducers. Values for different replicates separated with commas.

|  | Control | CIP | TET | CIP-TET |
| --- | --- | --- | --- | --- |
| Gly | 7.09,7.35,7.48 | 7.6,7.56 | 6.59,6.68,7.28 | 7.06,7.14 |
| Glu | 7.2,7.2 | 7.7,7.96 | 7.08,7.10 | 6.93,6.85 |
| Glu-aa | 7.04,7.01 | 7.6,7.37,7.29 | 7.31,6.94 | 7.19,7.14 |

